# Correlation and Matching Representations of Binocular Disparity across the Human Visual Cortex

**DOI:** 10.1101/2024.08.10.607440

**Authors:** Bayu Gautama Wundari, Ichiro Fujita, Hiroshi Ban

## Abstract

Seeing three-dimensional objects requires multiple stages of representational transformation, beginning in the primary visual cortex (V1). Here, neurons compute binocular disparity from the left and right retinal inputs through a mechanism similar to local cross-correlation. However, correlation-based representation is ambiguous because it is sensitive to disparities in both similar and dissimilar features between the eyes. Along the visual pathways, the representation transforms to a cross-matching basis, eliminating responses to falsely matched disparities. We investigated this transformation across the human visual areas using functional magnetic resonance imaging (fMRI) and computational modeling. By fitting a linear weighted sum of cross-correlation and cross-matching model representations to the brain’s representational structure of disparity, we found that areas V1-V3 exhibited stronger cross-correlation components, V3A/B, V7, and hV4 were slightly inclined towards cross-matching, and hMT+ was strongly engaged in cross-matching. To explore the underlying mechanism, we identified a deep neural network optimized for estimating disparity in natural scenes that matched human depth judgment in the random-dot stereograms used in the fMRI experiments. Despite not being constrained to match fMRI data, the network units’ responses progressed from cross-correlation to cross-matching across layers. Activation maximization analysis on the network suggests that the transformation incorporates three phases, each emphasizing different aspects of binocular similarity and dissimilarity for depth extraction. Our findings suggest a systematic distribution of both components throughout the visual cortex, with cross-matching playing a greater role in areas anterior to V3, and that the transformation exploits responses to false matches rather than discarding them.

**Significant Statement:** Humans perceive the visual world in 3D by exploiting binocular disparity. To achieve this, the brain transforms neural representation from the cross-correlation of signals from both eyes into a cross-matching representation, filtering out responses to disparities from falsely matched features. The location and mechanism of this transformation in the human brain are unclear. Using fMRI, we demonstrated that both representations were systematically distributed across the visual cortex, with cross-matching exerting a stronger effect in cortical areas anterior to V3. A neural network optimized for disparity estimation in natural scenes replicated human depth judgment in various stereograms and exhibited a similar transformation. The transformation from correlation to matching representation may be driven by performance optimization for depth extraction in natural environments.

## Introduction

The stereo-perceptual system forms a unified 3D perception from two slightly different 2D retinal images. This binocular disparity arises because our eyes view the same scene from different angles. To construct a veridical 3D representation, the system must solve the correspondence problem: determining which elements in the two eyes originate from the common spatial source (Julesz, 1960). Stereo computation starts in the primary visual cortex (V1), where neurons encode binocular disparity through computation similar to local cross-correlation (binocular energy model, BEM; Ohzawa et al., 1990). V1 neuron responses carry ambiguity as they signal disparity from both matched and mismatched features between the two eyes (Cumming and Parker, 1997). Solving the correspondence problem requires identifying correct and false matches and eliminating responses to false ones (Marr and Poggio, 1976).

Binocular disparity processing involves extensive visual areas (Welchman 2016; Rosenberg et al., 2023). In macaque monkeys, V4 neurons, but not MT neurons, show reduced sensitivity to disparities in falsely matched features (Krug et al., 2004; Tanabe et al., 2004; Yoshioka et al., 2021). In contrast, neurons in higher areas of both the ventral and dorsal pathways, i.e., inferotemporal (IT) and anterior intraparietal (AIP) cortices, become completely insensitive to 3D shapes defined by falsely matched features, forming a cross-matching representation (Fig. 1b; Janssen et al., 2003; Theys et al., 2012). However, the distribution of these representations in the human visual cortex is inconclusive, with findings conflicting with other human and monkey studies. For example, Bridge and Parker (2007) reported reduced sensitivity to false matches in V4, consistent with monkey neurophysiology, whereas Preston et al. (2008) did not observe similar results. Both studies agree that hMT+ exhibits less sensitivity to false matches, contrasting with findings in monkey MT (Krug et al., 2004). Human psychophysics suggests that the brain exploits both correlation and matching representations for stereopsis, even without achieving the correspondence (Fujita and Doi, 2016; Read 2023). Therefore, the map and mechanism of these representations within the human visual hierarchy remain elusive.

**Figure 1.**
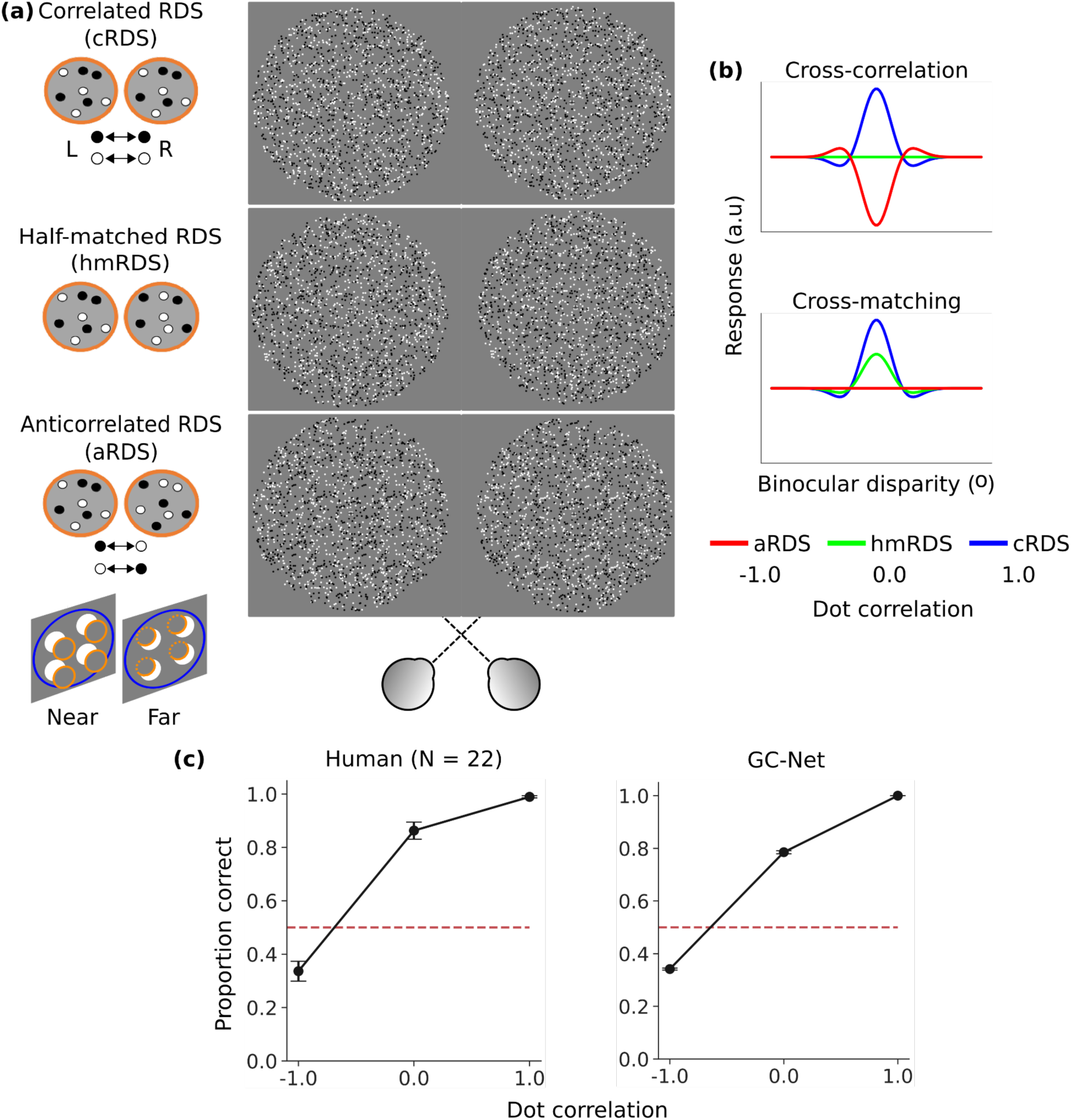
**(a)** Three RDS types were used: cRDS (binocular dot correlation = +1, comprising binocularly matching contrast dots), hmRDS (dot correlation = 0, blending contrast-matched and contrast–reversed dots equally), and aRDS (dot correlation = −1, displaying opposing contrast dots). A single type was presented identically in four circular RDSs (imaginary orange circles) surrounded by a cRDS background (imaginary blue circles). **(b)** Theoretical neural responses: Cross-correlation neurons are sensitive to disparity in cRDSs and aRDSs, with inverted tuning for aRDSs. Cross-matching neurons are sensitive to disparity in cRDSs and hmRDSs, but not in aRDSs. **(c)** Comparison of near/far judgment performance across the three RDS types between human observers (n = 22) and GC-Net (n=50, bootstrap). This comparison aimed to solely validate GC-Net for our simulation, not a main focus of this study (see Wundari and Ban, 2024 for the analysis). Briefly, both groups showed depth reversal for aRDSs (crossed disparity perceived as far and uncrossed disparity perceived as near) and “normal” depth for hmRDSs and cRDSs (cross disparity perceived as near and uncrossed disparity perceived as far). The error bars indicate the standard error across participants (human psychophysics) and bootstrap standard error (GC-Net).

We investigated these issues by measuring human visual cortex activity using fMRI. To evoke correlation-based and matching-based responses, we used random-dot stereograms (RDSs) with three dot correlation levels: correlated (cRDSs), anticorrelated (aRDSs), and half-matched (hmRDSs, Fig. 1a). cRDSs comprise dots with the same contrast polarity in both eyes, while aRDSs invert the polarity of corresponding dots. Neurons engaged in cross-correlation respond to aRDSs with inverted disparity tuning (Cumming and Parker, 1997), while those representing the solution to the correspondence problem are insensitive to disparity in aRDSs (Janssen et al., 2003; Theys et al., 2012). aRDSs induce the reversal of perceived depth (Tanabe et al., 2008; Zhaoping and Ackerman, 2018). hmRDSs, which blend contrast-matched and contrast-reversed dots equally, selectively activate cross-matching-based, not cross-correlation-based neurons, according to the BEM. The stimuli robustly elicit veridical depth sensation in humans (Doi et al., 2011, 2013). The use of hmRDSs advances previous fMRI studies (Bridge and Parker 2007; Preston et al., 2008) by providing a clearer dissociation between cross-correlation-based and cross-matching-based responses.

We estimated the contribution of each component to regional brain responses by comparing the similarity structure of brain and model disparity representations based on these computations. The analysis demonstrated that early areas V1-V3 primarily represent disparity through cross-correlation. Cross-matching slightly dominated later in V3A/B, V7 (IPS-0), and hV4, and was strongly represented in hMT+. To gain insights into the transformation, we simulated stereo processing using a deep neural network that matched human performance in depth judgment with cRDSs, hmRDSs, and aRDSs. Although the network was not constrained to match fMRI data, its disparity representation progressed from cross-correlation to cross-matching across layers, suggesting that optimization for depth extraction drives the transformation. Activation maximization analysis suggests that the transformation involves initial processing of extracting disparity from binocularly similar and dissimilar features, a middle phase focusing on processing of dissimilar features, and a final phase emphasizing matching processing.

## Materials and Methods

### Experimental Design and Statistical Analyses

We conducted fMRI measurements of brain responses while participants passively viewed cRDSs, hmRDSs, and aRDSs. We applied standard univariate and multivariate analyses, as well as model-based analyses, to analyze the fMRI data. We then simulated stereo processing with a deep neural network trained for depth estimation and analyzed its units to gain insights into the transformation process from cross-correlation to cross-matching representations. Detailed description of each experiment and analysis are provided below.

### Participants

Twenty-five participants [20 males and five females, 24.5 ± 4.37 (mean ± SD) years old] from Osaka University participated in the fMRI experiments. Three male participants were excluded from the data analysis: two did not complete the psychophysics experiment, and one had severe head movements during the fMRI sessions. They all had normal or corrected-to-normal vision. All procedures adhered to the Declaration of Helsinki 2008 and were approved by the Ethics and Safety Committees at the Center for Information and Neural Networks (CiNet), National Institute of Information and Communications Technology (NICT), and Osaka University. Participants provided written and oral informed consent before the experiment.

### Task and visual stimuli

Visual stimuli were generated and controlled using MATLAB (Mathworks Inc.) with psychtoolbox (Kleiner et al., 2007). The stimuli were back-projected onto a customized translucent screen (19.7° horizontally × 15.8° vertically in visual angle; Kiyohara Optics, Inc., Tokyo, Japan) behind the MRI bore. The stereo presentation was achieved using two LCD projectors (1920 × 1200 pixels, 60 Hz refresh rate; WUX4000, Canon, Tokyo, Japan), each with a distinct polarizing filter for the left and right images. The two projectors’ luminance values were gamma-corrected and matched using a MATLAB toolbox publicly available at https://github.com/hiroshiban/Mcalibrator2 (Ban and Yamamoto, 2013) and a colorimeter (Konica-Minolta, CS-100A). The filtered left and right RDS images were carefully aligned for perfect overlay on the screen. This was done by, prior to the experiment, aligning left and right images containing rectangle, circle, and grid patterns with different colors (https://github.com/hiroshiban/BinocularDisplayChecker). Participants viewed stimulus images through a tilted front-surface mirror, 96 cm away from the screen, wearing polarized glasses (KG-A04-B, Kiyohara Optics, Inc., Tokyo, Japan). Participants focused on a central fixation marker, a white square (0.6° visual angle), while performing a detection task to maintain attention (Popple et al., 1998). This task involved identifying the position of a randomly appearing white vertical bar displayed within the square for 250 ms. Participants pressed the left or right button depending on the bar’s position relative to the upper nonius line or refrained from responding if no bar appeared.

During the detection task, participants viewed dynamic RDSs (dot density = 25%, dot refresh rate = 30 Hz, painted on a mid-gray background; Tanabe et al., 2008). The timing of the stimulus presentation is detailed in the section on fMRI stereo experiments. The RDSs comprised 50% black and 50% white dots (0.14° diameter). Luminance levels were set at 128 cd/m^2^ for white dots, 64 cd/m^2^ for the mid-gray background, and 0.39 cd/m^2^ for black dots. The RDSs had a 14° (diameter) circular shape, with four identical smaller 4° (diameter) circular RDSs in each quadrant, centered at (2.5°, 2.5°) from the center of the background RDS (Fig. 1a). These four target RDSs were presented with disparities of −0.2° (crossed) or +0.2° (uncrossed). The background RDS, placed on the fixation plane, had a 0° disparity and was always binocularly correlated (i.e., black and white dots on the left image were consistently paired with black and white dots on the right image, respectively). The target RDSs displayed different levels of dot correlation: +1 (cRDSs), 0 (hmRDSs), and −1 (aRDSs). cRDSs comprise dots with the same contrast between the left and right images; aRDSs had opposing contrast dots—black dots in the left image paired with white dots in the right image, and vice versa—, and hmRDSs comprise an equal proportion of anticorrelated and correlated dots, thus displaying a net dot correlation of zero.

### fMRI image acquisition

The blood-oxygenation-level-dependent (BOLD) responses to stimuli were recorded using a Siemens MAGNETON Trio, A Tim System 3T Scanner at the NICT, CiNet imaging facility (Suita, Osaka, Japan) with a 32-channel phased-array head coil. Since we were only interested in the occipito-parietal cortex and participants wore polarized glasses during the recording, only the occipital part of the head coil was used. Voxel size was 2 mm isotropic, with a repetition time of 2 s, 78 slices, a multi-band factor of 3, a field of view of 192 × 192 mm^2^, and a flip angle of 75°. We collected 208 volumes for each functional run. The Multi-band EPI sequence was provided by the University of Minnesota (under a C2P contract, https://www.cmrr.umn.edu/multiband/).

High-resolution T1-weighted anatomical images of the whole brain (1 mm^3^ voxel resolution scan, 208 slices, TR = 1900 ms, echo time = 2.48 ms, field of view = 256 × 256 mm^2^, flip angle = 9°) were acquired for each participant for cortical surface reconstruction and precise coregistration between functional and structural images. We performed a separate session for localizing regions of interest.

### Retinotopic mapping and cortical preprocessing

Regions of interest (ROIs) were delineated for each participant using established retinotopic mapping procedures (Sereno et al., 1995; DeYoe et al., 1996; Dumoulin and Wandell, 2008; Yamamoto et al., 2008). These procedures involved identifying borders of retinotopic regions V1, V2, V3, V3A, V3B, V7, hV4, and hMT+ using rotating wedge stimuli and expanding/contracting concentric rings. Additionally, hMT+ boundaries for 12 out of the 22 participants were determined using hMT+ localizer stimulus, i.e., alternating motion and stationary dot patterns (Huk et al., 2002). Stimulus presentation codes are publicly available at https://github.com/hiroshiban/Retinotopy.

Cortical reconstruction was performed with BrainVoyager QX ver 2.8 (BrainInnovation B.V, Maastricht, The Netherlands), following established procedures (Ban et al., 2012). This process included stripping away the brain’s non-cortical structure (skull), transforming anatomical scans into Talairach space, inflating the cortex, and creating a flattened surface of the two cerebral hemispheres.

### fMRI stereo experiments

Imaging sessions employed a block design paradigm. Each functional run comprised 192 trials, starting and ending with a 16 s stimulus-off (mid-gray background with a fixation marker). One block lasted 16 s, with 1 s stimulus presentation periods alternating with 1 s stimulus-off periods (background only). The total duration per functional scan run was 416 s (i.e., 16 s stimulus-off + 24 blocks × 8 trials-per-block × 2 s-per-trial + 16 s stimulus-off). Within each block, only one of the six stimuli (cRDS, hmRDS, and aRDS, each with crossed or uncrossed disparity) was presented. The different stimuli were presented in a randomized order across the blocks. A 10 s “dummy” scan was inserted at the beginning of each scan to avoid transient signal drifts.

### fMRI data preprocessing

We preprocessed the MRI functional data using BrainVoyager QX ver 2.8. The first five volumes of each functional run were excluded to mitigate the transient magnetization effects common at the start of imaging sessions. The preprocessing pipeline included slice time correction, 3D motion correction, linear trend removal, and high-pass temporal filtering with a cutoff of three cycles per run. We omitted spatial smoothing to avoid information loss. The processed data were aligned with each participant’s anatomical scan and converted into Talairach space. Data from different runs were aligned to the initial volume of the first run for consistent orientation and scale across different runs.

### Univariate and multivariate analysis

We performed univariate and multivariate analyses to determine whether our fMRI data contained information about the experimental conditions. To select voxels used for these analyses, we employed a general linear model (GLM) in assessing the contribution of each experimental condition to the voxel’s time course variance. We were interested in whether the effect size contributed by all stimulus conditions was more significant than the baseline condition. The analysis was done for each voxel in the left and right hemispheres by incorporating the experimental conditions (3 RDS types x 2 disparity signs), six motion parameters (head movements: translations in XYZ axes and rotations in pitch, roll, and yaw), and a constant term as regressors. The regression coefficients (GLM beta values) were derived from the least-squares solution to the linear equation. We then computed the *t*-statistic for each voxel within the predefined brain regions with a contrast vector fixation *vs.* all conditions. The top 250 task-sensitive voxels were selected based on descending *t*-values.

For univariate analysis, we computed percent BOLD signal change from fixation baseline using voxels selected based on GLM analysis. The percent signal change was calculated as the average between crossed and uncrossed responses for each RDS type because we primarily focused on the effects of dot-correlation levels on overall regional activation.

For multivariate analysis or multi-voxel pattern analysis (MVPA), we used a linear support vector machine (SVM) to decode brain activity patterns associated with crossed and uncrossed disparities in RDSs, using the 250 most task-sensitive voxels as the feature dimension. This analysis was implemented via the sklearn library in Python (https://scikit-learn.org/stable/), with a linear kernel, the regularization parameter *c* was set to 1 to give a hard margin, and default settings were used for other parameters. For each participant, we cross-validated the fMRI dataset for each ROI and RDS type with a leave-one-run-out scheme to evaluate the decoder’s generalization ability. The final decoding accuracy for each RDS type was averaged across participants.

The statistical significance (*p*-value) of the classifier’s mean prediction accuracy for the selected voxels in each ROI was assessed against the chance performance baseline. Multivariate normality assumed in the classical multivariate methods is often violated in fMRI data (Kriegeskorte, 2011). To ensure the statistical reliability of our MVPA results without relying on this assumption, we conducted a permutation test and bootstrap resampling for the group-level analysis across participants. The permutation test establishes a baseline for chance performance, while bootstrap resampling estimates a *p*-value for MVPA against a null distribution to which the true accuracy (obtained from the data) can be compared. The null distribution reflects the decoding accuracies computed when randomizing the task labels and the voxel values disrupted the relationship between the task labels (Etzel, 2015). These tests require no distributional assumptions but are computationally expensive (Nichols and Holmes, 2002).

For the permutation test, we generated a null distribution for each RDS type: cRDS, hmRDS, and aRDS. We shuffled the task labels (near and far) to the individual’s BOLD patterns dataset for each ROI and RDS type, then performed a decoding analysis using a leave-one-run-out cross-validation scheme. This process was repeated for all participants. The decoding accuracies for each RDS type were then averaged across participants. We iterated the same process 10,000 times to generate a null distribution of 10,000 classification accuracies per RDS type. The baseline of chance performance level for each RDS type was defined as the 95^th^ percentile of the corresponding null distribution.

We generated a bootstrap distribution by resampling, with replacement, individual decoding performance for each ROI and RDS type. The mean of the resampled data was calculated, and this step was repeated 10,000 times. We estimated the *p*-value for each ROI and RDS type by calculating the proportion of the resulting 10,000 classification accuracies below the baseline computed in the permutation test. Following a two-tailed test and adjusting for multiple comparisons across ROIs, the confidence level alpha was set at 0.003125 [alpha = 0.05/(2 × 8 (ROIs)) = 0.003125]. A *p*-value below this threshold indicated statistical significance.

### Searchlight analysis

We performed the searchlight analysis (Kriegeskorte et al., 2006) to ensure our predefined ROIs using a separate retinotopic scanning covering the essential brain regions associated with the experimental conditions. We computed the average of local multivariate searchlight across participants to generate an information-based map containing brain activity patterns representing crossed- and uncrossed-disparity information in cRDSs. We then used a 6 mm (radius) sphere to decode locally (using SVM) the brain activity patterns throughout the whole cortex in the Talairach space. The resulting searchlight map was overlaid to a representative individual’s brain.

### A linear combination of cross-correlation and cross-matching encoding models

We hypothesized that the brain encodes two disparity representations derived from the left and right images: cross-correlation and cross-matching. We proposed a model which computes a linear weighted sum of cross-correlation and cross-matching representations (Correlation and Matching Model, CMM) to study the transformation of disparity information across visual cortices. The CMM posits that disparity representation in each cortical area can be approximated as a linear weighted sum of these two representations. The linear model successfully explains the psychometric function of perceptual depth in human observers elicited by aRDSs, hmRDSs, and cRDSs (Tanabe et al., 2008; Doi et al., 2011, 2013; Fujita and Doi, 2016). It is important to note that we did not impose any representational transformation in the CMM modeling.

Neural activity based on a cross-correlation computation linearly correlates with the correlation level between the left and right retinal images, responding to both correlated (cRDSs) and anticorrelated (aRDSs) features, resulting in ambiguous disparity signal (Cumming and Parker, 1997; Read and Cumming, 2007; Goncalves and Welchman, 2017). These neurons exhibit a flat disparity tuning curve for hmRDSs, which have zero interocular correlation (upper panel of Fig. 1b). In contrast, cross-matching neurons display non-linearity as they filter out responses to negatively correlated features. They only fire when the correlation between the left and right inputs is greater than zero (Fig. 1b, lower panel), making them insensitive to false matches and focused on correct global matches (Nieder & Wagner, 2000; Janssen et al., 2003; Tanabe et al., 2004; Abdolrahmani et al., 2016; Yoshioka et al., 2021). Psychophysical evidence suggests that humans rely on both neural representations to infer depth from RDSs at various dot correlation levels (Tanabe et al., 2008; Doi et al., 2011, 2013; Aoki et al., 2017; for a review, see Fujita and Doi, 2016).

To compare the brain and CMM representations, we performed the modeling within the framework of representational similarity analysis (RSA, Kriegeskorte et al., 2008). We computed representational dissimilarity matrices, RDMs, for empirical fMRI data and simulated data from the CMM. An RDM is a matrix that captures all possible pairwise distances in the representational space among response patterns (Kriegeskorte et al., 2008; Kriegeskorte and Diedrichsen, 2016; Kriegeskorte and Wei, 2021). RDMs were generated by comparing the activity patterns associated with each pair of experimental conditions: cRDS- crossed, cRDS-uncrossed, hmRDS-crossed, hmRDS-uncrossed, aRDS-crossed, and aRDS- uncrossed (Fig. 2a, left panel).

**Figure 2.**
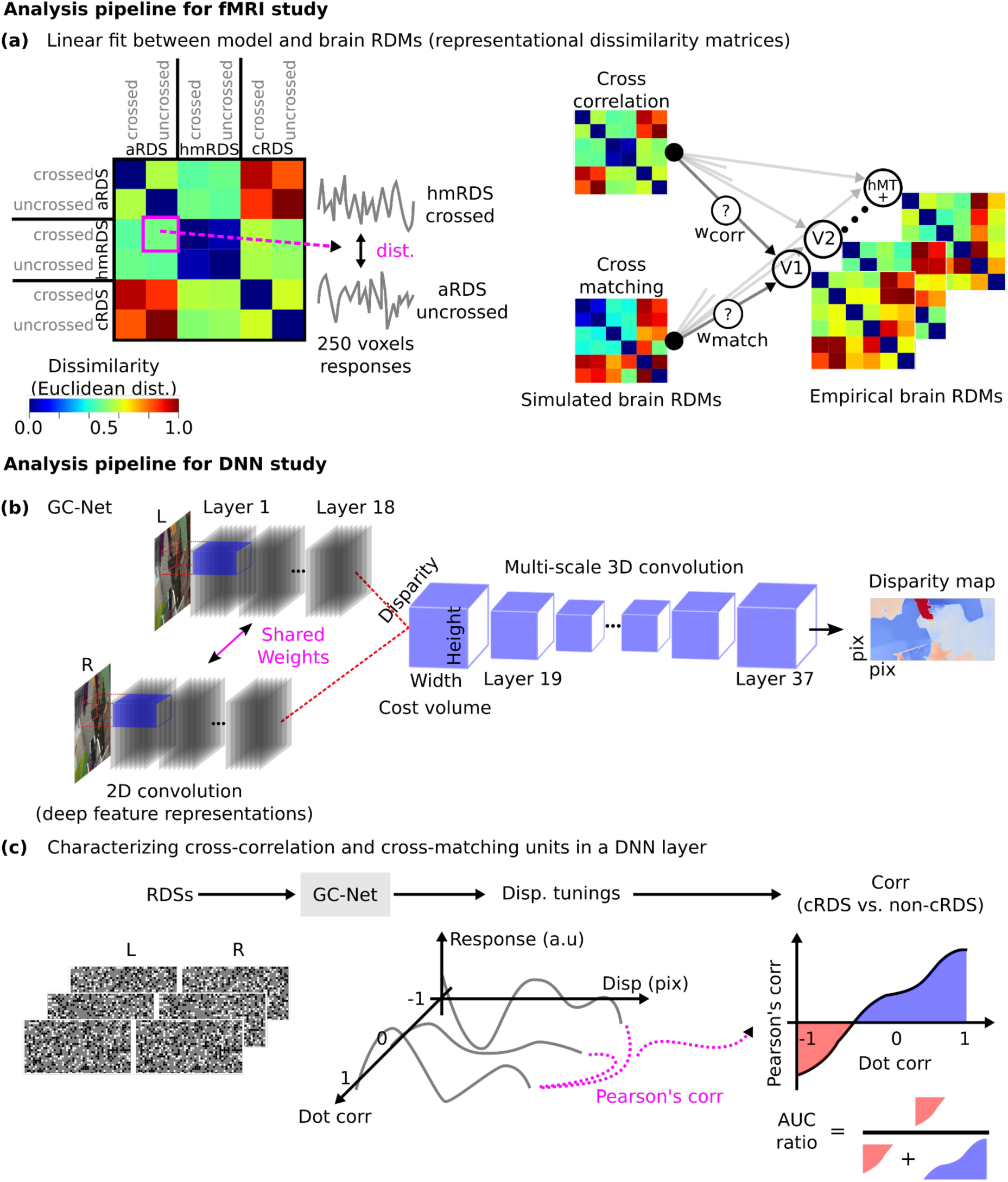
Analysis pipelines for fMRI **(a)** and DNN **(b, c)** studies. **(a)** Left: Stimuli-pairwise configurations in our representational dissimilarity matrix (RDM). Each element represents the normalized Euclidean distance between response patterns to RDSs. Right: Model and empirical brain RDMs, with model RDMs used as regressors for empirical brain RDMs. Connecting lines indicate regressor weights. **(b)** Model architecture of the Geometry and Context Network (GC-Net; Kendall et al., 2017). **(c)** Characterization of GC-Net units’ computational types using RDSs with varying dot correlations and disparities. The area under the curve (AUC) ratio was used to determine the preference for cross-correlation or cross-matching.

### Generating the RDMs for cross-correlation and cross-matching models

We computed two model RDMs, one for cross-correlation and one for cross-matching (Fig. 2a, left panel). The (i, j) element in an RDM denotes the Euclidean distance between 250 simulated voxel responses for experimental conditions (i) and (j). We simulated six conditions: cRDS-crossed, cRDS-uncrossed, hmRDS-crossed, hmRDS-uncrossed, aRDS-crossed, and aRDS- uncrossed. We simulated 500 RDSs per type (120 x 120 pixels) with disparities of ± 10 pixels (1 pixel = 0.02°). This process was repeated 1,000 times.

Each 2 mm voxel mimicked a ‘disparity column map’ reflecting the visual cortex’s columnar organization for disparity neurons (DeAngelis and Newsome, 1999; Adams and Zeki, 2001; Yoshiyama et al., 2004; Tanabe et al., 2005; Chen et al., 2008). This map was ordered by disparity preference along the cortex’s tangential dimension, modeled by a sawtooth wave profile with 3 mm/cycle, simulating the gradual and systematic change observed in the cortical columns (Kamitani and Tong, 2005; Ban et al., 2012). Disparity preferences ranged from −0.5° to 0.5° in 0.05° increments. We added Gaussian noise (centered at 0) to the sawtooth profile for variability in disparity preference within each voxel.

The synthetic disparity column map grouped units within a voxel size of 2 mm. Each voxel comprised disparity-selective units tuned to spatial frequencies of 1, 2, 4, 8, and 16 cycles/°. A voxel’s response was defined as the sum of all disparity-selective neuron responses across these frequencies.

We calculated the response of a disparity-sensitive unit using cross-correlation and cross-matching encoding algorithms. For cross-correlation, the response for a given disparity was derived from cross-correlating the left and right RDS images within a spatial window. The unit response to a single trial was the sum of pixel values within a square window without any weighting function (without a specific receptive field or RF structure). Formally, cross-correlation response *C* at disparity *d* is expressed as (Doi and Fujita, 2014):

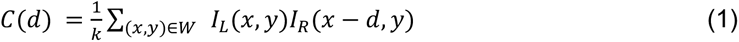

where *I_L_* and *I_R_* indicate the left and right RDSs, respectively; x and y represent the horizontal and vertical pixel coordinates, respectively; d indicates the horizontal disparity between the left and right spatial window W; and k indicates the number of elements (pixels) in the spatial window. The elements inside the RDS matrix only have three values: −1 (for black dots), 0 (for gray background), and +1 (for white dots). Thus, C(d) is assigned +1 for contrast-matched pixels (black-black or white-white), −1 for contrast-reversed pixels (black-white), and 0 for any pixel meeting the background pixels.

Cross-matching is similar to cross-correlation but includes a half-wave rectifier before averaging. The cross-matching response *M* has the following form (Doi and Fujita, 2014):

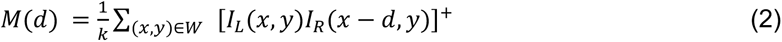

where [.]^+^ indicates half-wave rectification. This nonlinear operation takes place at the pixel level. Thus, 0 will be assigned for every contrast-reversed pixel.

The size of the spatial square window *W* in eq. 1 and 2 was the full-width half-maximum (FWHM) bandwidth of around 1.5 octaves, inversely proportional to the units’ preferred spatial frequency (1, 2, 4, 8, and 16 cycles/°) in the simulated voxel, as defined in eq. 3 (Read and Cumming, 2007):

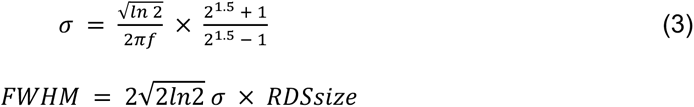

### Fitting CMM RDMs to fMRI RDMs

Figure 2a outlines the analysis pipeline for our fMRI study. We consider that disparity representation in each visual area cannot be explained by a single feature set, but by a mixture of different representations. A simple approach to implement this idea is by fitting a regional brain RDM using a linear weighted sum of the cross-correlation and cross-matching RDMs. A linear combination of feature representations using RDM has been used to explain object representations in IT (Khaligh-Razavi and Kriegeskorte, 2014).

We took the upper triangular halves of the empirical and model RDMs as vectors, mean-centered, and max-normalized for scale consistency. We implemented nonnegative least squares with the L2-norm (Ridge regression, through scikit toolbox, https://scikit-learn.org/stable/, with regularization parameter alpha = 1 (default), solver = “lbfgs”, positive = True) for the fitting. The nonnegativity ensured positive weights for interpretability, and L2-norm was used to mitigate the multicollinearity effect (the problem in regression in which predictor variables are correlated) between the two regressors (the cross-correlation and cross-matching RDMs). We performed leave-one-subject-out cross-validation and bootstrapped over 1,000 iterations to mitigate overfitting. The resulting weights indicate the proportions of cross-correlation and cross-matching components contributing to disparity representation. Comparing an empirical RDM from a given ROI with model RDMs evaluates the similarity of disparity representation of that ROI to the combination of cross-correlation and cross-matching representations. This allows us to trace the representational transformation throughout the visual cortex.

### Calculating the goodness of fit of the model and noise ceiling

We assessed the fit of our model RDM to the brain RDM using the Kendall rank correlation coefficient (the Kendall-*τ* type b from the SciPy library in Python). Kendall-𝛕_B_ measures the similarity between two data after ranking the quantities and returns a value between 0 (no correlation) and 1 (perfect correlation).

Noise in the fMRI data can make estimating the performance of the true model uncertain. Therefore, we estimated a noise ceiling to define the lower and upper limits that the true model can achieve (Nili et al., 2014). For each ROI, we estimated the upper bound by first averaging the RDM across individuals and assigning it as a reference RDM. We randomly sampled RDMs (22 individuals) with replacement and averaged the sampled RDMs to get a candidate RDM. We bootstrapped this process 1000 times, resulting in a bootstrap population containing 1000 candidate RDMs. We calculated the Kendall-𝛕_B_ rank correlation between the reference RDM and each candidate RDM in the bootstrap population and took the median of the 1000 Kendall-𝛕_B_ population as the upper bound. A similar approach was used for the lower bound; however, we used an individual RDM as the reference RDM. To this end, we selected one individual RDM and performed 1,000 bootstrap iterations with the remaining 21 RDMs to get a population of candidate RDMs. We calculated the Kendall-𝛕_B_ correlation between the reference RDM and each candidate RDM in the bootstrap population, and we took the median of the Kendall-𝛕_B_ population. The process was repeated until all individual RDMs were assigned as reference RDMs. Lastly, we averaged these median values to determine the lower bound.

### Simulation of disparity processing with a deep neural network

According to the BEM, disparity information is encoded by the interocular correlation between the left and right RFs (Ohzawa et al., 1990, Read et al., 2002). This drives us to examine disparity signals modulated by cross-correlation and cross-matching neurons across the visual hierarchy. However, given the impossibility of precisely identifying cross-correlation and cross-matching neurons in fMRI voxels, we explored the question by simulating disparity processing in a deep neural network (DNN) optimized for processing binocular disparity. DNNs share similarities in processing visual information with human and non-human primate brains as the result from optimization for a given task (Khaligh-Razavi and Kriegeskorte, 2014; Yamins et al., 2014; Güçlü and van Gerven, 2015; Kriegeskorte, 2015; Yamins and DiCarlo, 2016).

Based on this reasoning, we explored the stereo coding underlying the transformation from cross-correlation to cross-matching using the Geometry and Context Network (GC-Net; Kendall et al., 2017). GC-Net incorporates contextual information, such as details about the surroundings of the object of interest, to estimate horizontal binocular disparity from pairs of rectified (vertically aligned) stereo images. We implemented GC-Net (Fig. 2b) using PyTorch 2.1 (Paszke et al., 2019) and trained it on the Scene Flow datasets (https://lmb.informatik.uni-freiburg.de/resources/datasets/SceneFlowDatasets.en.html; Mayer et al., 2016). The datasets comprise three subsets, each with a different contextual environment: FlyingThings3D, Monkaa, and Driving.

The details on the layer dimensions of the GC-Net can be found in the original paper (Kendall et al., 2017). Briefly, GC-Net comprised 37 layers, with the first 18 layers forming convolutional layers to extract deep feature representations from left and right image inputs. The outputs from these left and right channels in layer 18 were concatenated at the cost volume layer corresponding to each disparity level (total pixel displacement D = 192 pixels, from −96 to 95 pixels with increments of 1 pixel), resulting in a 4-dimensional tensor for each stereo image. The representation in the cost volume was refined through multiple stages of 3-dimensional convolutions in a U-Net structure (deep encoder-decoder) to estimate disparity. Signals from the left and right channels first merged and interacted in layer 19.

The code of GC-Net was adapted from an open-source implementation (https://github.com/zyf12389/GC-Net/tree/master) with modifications in the creation of cost volume tensor. For training, we implemented the AdamW optimizer in PyTorch (Loshchilov and Huttler, 2017) with a learning rate of 6 × 10^−4^ and a learning rate decay of 0.2 per epoch. We used a batch size of two due to the limit of our GPU memory and ten epochs (one epoch is a complete cycle through the training data). The image dataset was split into 80% for training and 20% for validation. The original images (540 x 960 pixels) were cropped to a patch size of 256 x 512 pixels, with the cropping location randomized across the entire images. This ensured the network received different parts of the images in each epoch.

We used the L1 loss, the mean absolute error between the predicted disparity map and the ground truth, to assess the network’s performance. The loss was calculated every 200 iterations and averaged over 25 iterations to ensure stability and reliability in our metrics. We selected the model’s parameters with the lowest L1 loss on the validation dataset. The training was accelerated on a GPU NVIDIA RTX 4090 (Nvidia, California, USA).

The formation of the cost volume tensor (after layer 18) was modified from Kendall et al. (2017) to enable GC-Net to predict disparity from (−D/2 = −96 pixels) to (+D/2 - 1 = 95 pixels), instead from 0 to 192 pixels. For each stereo image, we subtracted the true disparity map from 1.5 times its median value, allowing the disparity to have negative values. After multiple stages of 2D convolutions up to layer 18, the size of the left and right input tensors became [B, F, H/2, W/2], where B = batch size, F = feature size, H = image height, and W = image width. For each stereo image, we concatenated each right input feature with its corresponding left input feature for each disparity level. The concatenation was done by padding the right input features in layer 18 with zeros on their left and right sides (the padded tensor size became [B, F, H/2, (D/4 | W/2 | D/4)]). For the left input features in layer 18, the padding was done iteratively from 0 to D/2 (the padded tensor size became [B, F, H/2, (d_i | W/2 | D/2 - d_i)], for d_i in range from 0 to D/2). In each iteration, the padded right feature was concatenated with the padded left feature along the feature axis and appended to the disparity axis, resulting in a dimension [B, D, 2F, H, W/2 + D/2].

### Characterizing cross-correlation and cross-matching units in GC-Net

After training GC-Net, we characterized the processing units in each layer to determine whether they were involved in cross-correlation or cross-matching (Fig. 2c). We fed the network with RDSs varying in dot correlation (ranging from −1.0 to +1.0, in increments of 0.2, corresponding to dot match ranging from 0.0 to +1.0 in increments of 0.1) and disparity magnitudes from −30 to +30 pixels (in increments of 2 pixels), producing disparity tuning for each unit across dot correlation levels in each layer.

Next, we calculated Pearson’s correlation between the disparity tuning for cRDS (dot correlation of +1.0) and non-cRDS (dot correlation from −1.0 to +0.8). This resulted in a correlation function across dot correlation levels. The preference for cross-correlation or cross-matching was quantified using the area under the curve (AUC) ratio of this correlation function (Fig. 2c, right panel; “area ratio” defined in Yoshioka et al., 2021). This metric compares the AUC for correlations below zero (shaded in red) and the AUC for correlations above zero (shaded in blue). A unit with a pure cross-matching computation should have an AUC ratio of zero, because Pearson’s correlation does not become negative across dot- correlation levels. A unit with a pure cross-correlation computation should have a ratio of one, because this unit will invert tuning functions other than cRDSs.

We did not use the amplitude ratio or signed amplitude ratio based on Gabor fit to determine the preference for cross-correlation or cross-matching, as defined in previous studies (Cumming and Parker, 1997; Yoshioka et al., 2021). We reasoned that using peak amplitude from Gabor fitting overlooks other essential properties of disparity tuning, such as the width and sidelobes of the tuning function, which convey important information about the disparity. Instead, we used Pearson’s correlation as it accommodates the amplitude, width, and sidelobes of the tuning. The sign of the correlation coefficient indicates whether the tuning inversion occurs, while its magnitude evaluates the strength of the unit’s preference for cross-correlation or cross-matching.

### Activation maximization

We characterized the left and right preferred inputs of each unit in the GC-Net to understand how monocular receptive field structures evolve across layers to support stereo processing. We employed activation maximization technique or feature visualization via optimization (Erhan et al., 2009; Zeiler and Fergus, 2014; Olah et al., 2020a, b; Voss et al., 2021), which uses gradient ascent to find the inputs that maximize the activation of units or parts of the network. We then calculated Pearson’s correlation between the left and right preferred inputs (receptive fields) to measure their spatial structure similarity. This approach is essential to understand how units at each stage process disparity information.

Our activation maximization analysis was facilitated by the Captum library (https://captum.ai/), a versatile tool for interpretability in neural networks. We used the “neuron objective” (the loss that maximizes the activations of a target unit in a specified channel from a specified layer) at the center of the layer in each of the feature and disparity channels in layers 19–37. We started the analyses from layer 19, where signals from the left and right channels first merge and interact. We maximized the pre-activation value (the convolutional layer output) of the central unit in each feature and disparity channel feeding into the softmax layer.

### Data and code availability

Python codes for this article are available at https://github.com/wundari/CMM_model. Preprocessed fMRI data, pretrained model, and processed files are available upon request. This material has not been peer reviewed.

## Results

Participants first underwent standard retinotopic mapping of the visual cortex using fMRI, viewing rotating wedges and expanding/contracting annuli, to delineate the borders of visual areas V1, V2, V3, V3A, V3B, V7, hV4, and hMT+ (Fig. 3). We then performed a searchlight analysis (Kriegeskorte et al., 2006) to verify that our predefined ROIs covered the essential brain regions involved in the experimental conditions. The resulting searchlight map indicated that multiple visual areas in both hemispheres represent the near/far plane defined in cRDSs (Fig. 3). Our predefined ROIs encapsulated most regions where local activity patterns are associated with processing disparity information, except for the region dorsal to V7, likely corresponding to IPS1/2/3 (Swisher et al., 2007). We confined our analysis to these retinotopic areas identified separately from the searchlight map using passively viewed rotating wedges and expanding/contracting annuli (dashed black line in Fig. 3).

**Figure 3.**
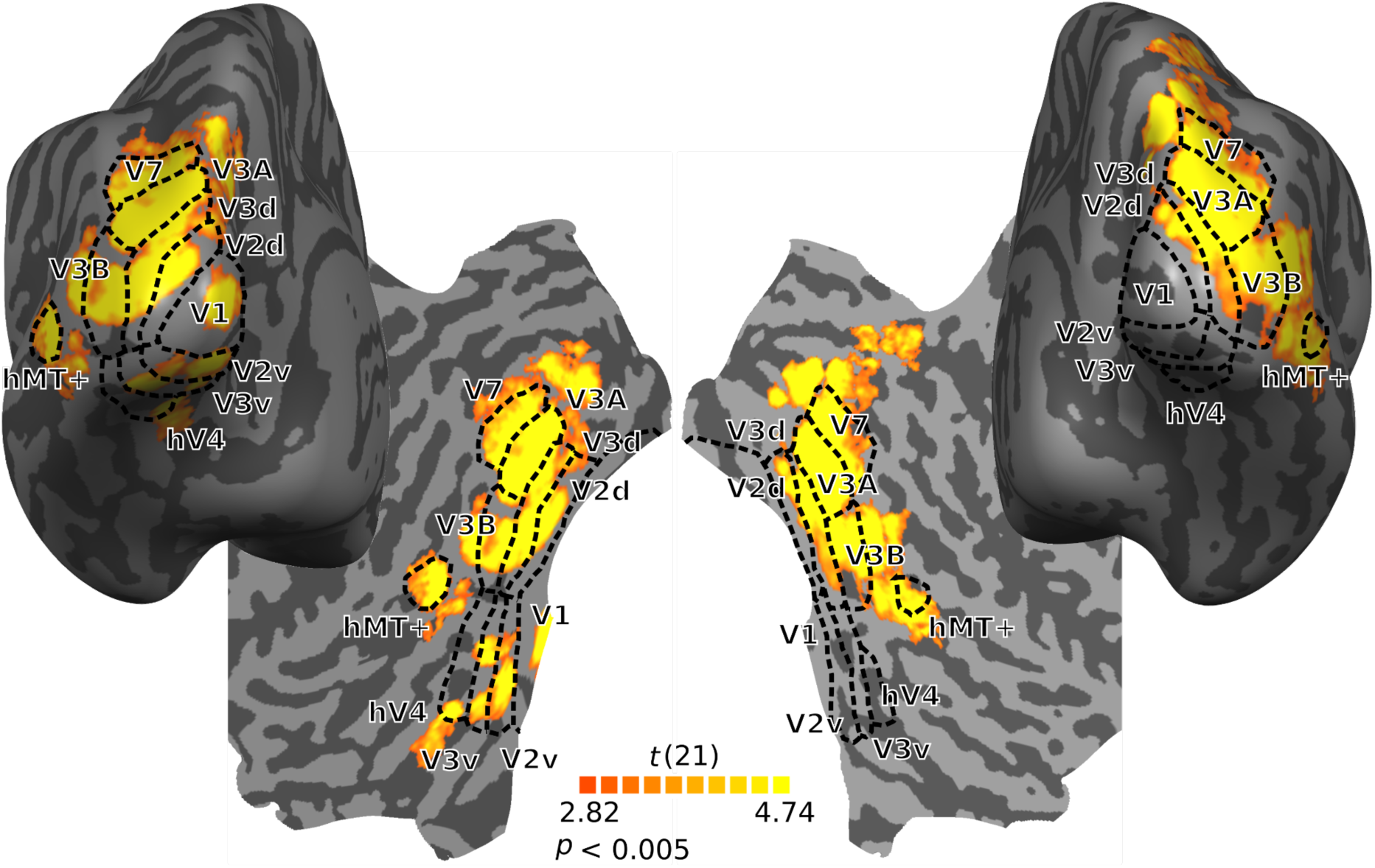
The searchlight map overlaid on the inflated and flattened left and right hemispheres. Sulci appear in a darker gray than gyri. Dashed black lines demarcate the retinotopic area boundaries. Color coding indicates *t*-values from response pattern classification accuracies in a group searchlight analysis (Kriegeskorte et al., 2006), marking brain regions whose activity patterns carry information regarding disparity categories (crossed vs. uncrossed) in cRDSs.

### BOLD activation to dynamic RDSs with varying dot correlations across ROIs

We presented dynamic RDSs with three levels of binocular dot correlation (Fig. 1a; cRDSs, hmRDSs, and aRDSs) to evoke cross-correlation and cross-matching BOLD responses (see Materials and Methods). We performed a univariate analysis to examine whether regional BOLD responses to these RDSs exhibited distinct overall activities across brain regions. Although cross-correlation and cross-matching in principle should be reflected in aRDS and hmRDS brain activities, respectively, we performed the analysis on cRDSs as well, important as a control measure. We quantified brain region activation to these stimuli by calculating the percent signal change relative to the fixation-only condition (Fig. 4). A two-way ANOVA revealed a significant main effect of cortical area (*F_ROI_*(7, 504) = 169.78, *p* < 0.001), and dot correlation (*F_RDS_*(2, 504) = 11.40, *p* < 0.001). There was no significant interaction between ROI and RDS type (*F_RDSxROI_*(14, 504) = 0.17, *p* = 0.99).

**Figure 4.**
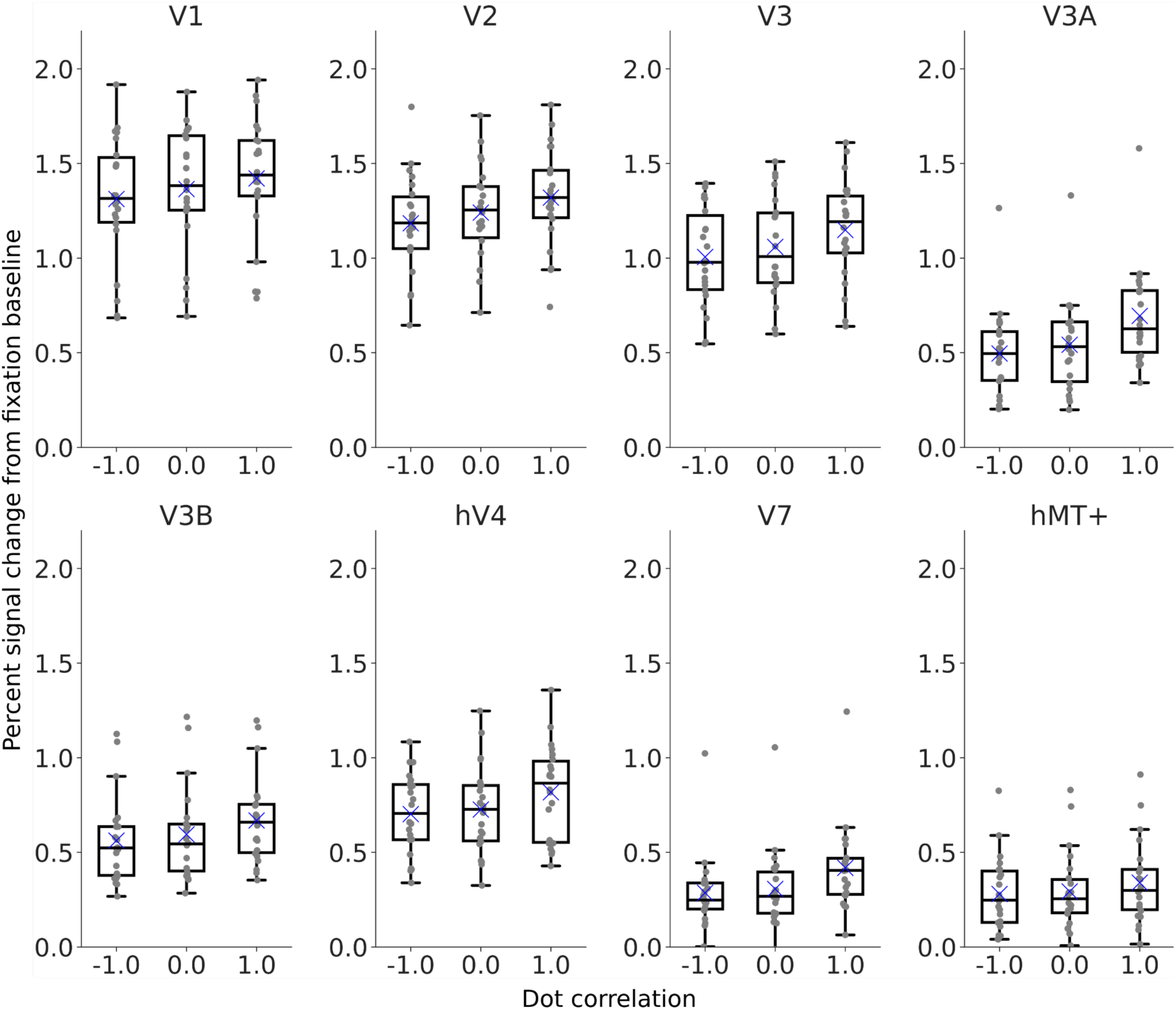
Box plots showing percent signal change relative to fixation baseline for the 250 most strongly modulated voxels within each predefined ROI. The box extends from the first quartile (Q1) to the third quartile (Q3) of the data, with a line at the median and a blue cross at the mean. The whiskers extend from the box to the farthest data point within 1.5 x the interquartile range (IQR). Gray dots represent individual participant data.

Pairwise comparisons using Mann-Whitney tests indicated differences in percent signal change between cRDSs and aRDSs in V3A (*p* = 0.0089) and V7 (*p* = 0.0058). However, these differences fell short of statistical significance when corrected for multiple comparisons (*p* = 0.003125), in agreement with Bridge & Parker (2007) but inconsistent with Preston et al. (2008) which found significantly stronger responses to cRDSs than aRDSs. Our results did not replicate the differential responses to cRDSs and aRDSs in hV4 and hMT+ reported in previous papers (Bridge and Parker 2007; Preston et al., 2008) (hV4: *p* = 0.14, hMT+: *p* = 0.39). When comparing between hmRDSs and cRDSs, we again found differences, but not significant, in V3A (*p* = 0.034) and V7 (*p* = 0.026). We did not detect any significant difference in activation between aRDSs and hmRDSs across all predefined ROIs, despite participants robustly perceiving veridical depth in hmRDSs and reversed depth in aRDSs (Fig. 1c). These results suggest that the average activation of individual areas at the group level cannot reliably dissociate cross-correlation from cross-matching responses, nor can they explain the distinct depth percepts elicited by cRDSs, hmRDSs, and aRDSs.

### Decoding disparity information across different RDS types and brain areas

The failure to detect differences in the mean responses among the three RDS types using the univariate approach prompted us to perform multivariate analysis. We conducted a decoding analysis on voxel response patterns in each ROI to assess whether information about crossed and uncrossed disparities in each RDS type could be extracted from spatial patterns of population activity (Fig. 5).

**Figure 5.**
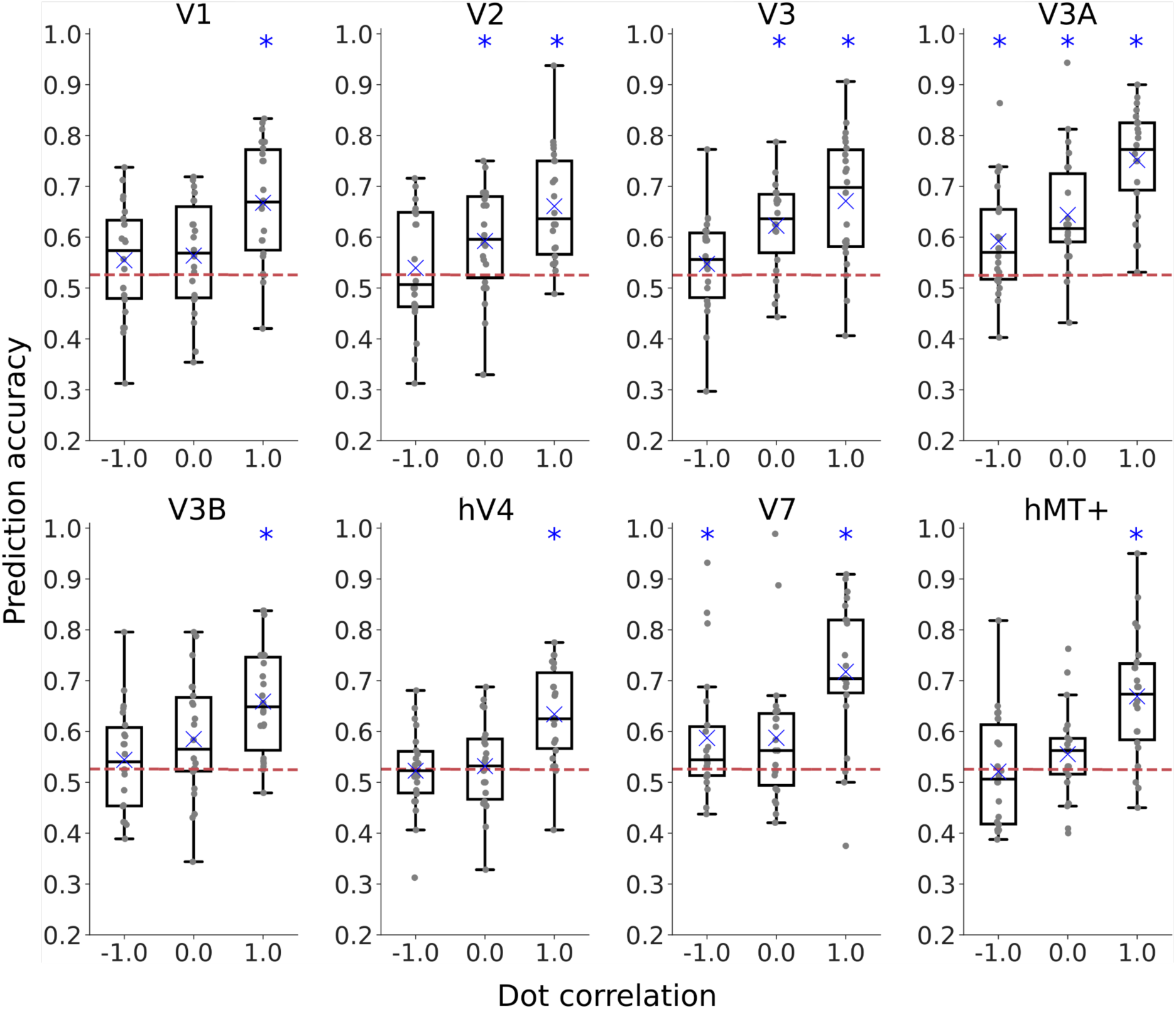
SVM decoding performance for discriminating between crossed and uncrossed disparities based on BOLD responses in predefined ROIs. Box plot conventions follow those in Figure 4. Gray dots represent individual participant data (*n* = 22). Red lines indicate the baseline chance performance from the 95^th^ percentile of 10,000 iterations in the permutation test for each dot correlation level. Asterisks indicate statistically significant decoding performance above the baseline, based on permutation test and bootstrap resampling [alpha = 0.05, *p* < 0.003125, adjusted for multiple comparisons across ROIs].

The prediction accuracies of the linear classifier (SVM) in discriminating between crossed and uncrossed disparities varied across RDS types and brain areas. The SVM successfully extracted stimulus disparity sign information from cRDS response patterns in all predefined ROIs: V1, V2, V3, V3A, V3B, hV4, V7, and hMT+, consistent with a previous study (Preston et al., 2008). The prediction accuracy for cRDSs ranged from 0.63 in hV4 to 0.75 in V3A, all of which was significantly above the chance performance baseline estimated by permutation test and bootstrap resampling (*p* < 0.0001).

We found that disparity signs in aRDSs and hmRDSs could only be decoded reliably from response patterns in V3A (aRDSs: mean accuracy = 0.59, *p* = 0.0004; hmRDSs: mean accuracy = 0.64, *p* < 0.0001). This suggests that V3A exploits the cross-correlation feature in aRDSs and the cross-matching feature in hmRDSs. Disparity sign information in aRDSs could also be reliably extracted from activity patterns in V7 (aRDSs: mean accuracy = 0.59, *p* = 0.0031), suggesting that cross-correlation was explicitly represented there. Reliable decoding performance for hmRDS was found in V2 (mean accuracy = 0.59, *p* = 0.0026) and V3 (mean accuracy = 0.62, *p* < 0.0001), suggesting explicit cross-matching representation in these early areas.

aRDSs decoding accuracy was increasingly robust from V1 (mean accuracy = 0.55, *p* = 0.11) to V3A (mean accuracy = 0.59, *p* = 0.0004). This result contradicts Preston et al. (2008), which reported attenuation of disparity information from V1 to V7 (with an initial increase from V1 to V2). Similarly, hmRDS decoding accuracy increased from V1 (mean accuracy = 0.56, *p* = 0.052) to V3A (mean accuracy = 0.64, *p* < 0.0001). Disparity information in aRDSs and hmRDSs attenuated in hV4 (aRDS: mean accuracy = 0.52, *p* = 0.58; hmRDS: mean accuracy = 0.53, *p* = 0.37).

The results of this multivariate analysis suggest that cross-correlation and cross-matching are present in the form of voxel fine-scale activity patterns spatially distributed across a cortical area.

### Proportion of cross-correlation and cross-matching across the visual hierarchy

Prior analyses verified the presence of cross-correlation and cross-matching representations in our fMRI data. However, the classifier in the multivariate analysis was not trained to match the brain responses *per se*, but rather to develop a decision rule that maximizes the classification accuracy in distinguishing BOLD response patterns associated with experimental conditions. While one might expect voxels informative about cross-correlation and cross-matching to carry stronger weights to decode aRDSs and hmRDSs, respectively, this assumption is not guaranteed. Other scenarios, such as weaker weights for those voxels or the involvement of voxels unrelated to either computational component, may also lead to successful decoding. This suggests that classification accuracy may not accurately reflect the proportion of cross-correlation and cross-matching components.

Therefore, we next attempted to quantify the contribution of cross-correlation and cross-matching to brain responses in each ROI by linearly combining the similarity structure of representation based on cross-correlation and cross-matching models (CMM) to fit the similarity structure of regional brain representation. The linearity assumption is reasonable as it successfully explains the psychometric function of perceptual depth in human observers evoked by aRDSs, hmRDSs, and cRDSs (Tanabe et al., 2008; Doi et al., 2011, 2013; Fujita and Doi, 2016). We performed the analysis in the representational similarity analysis (RSA) framework that examines the structure of disparity representations within a representational space based on relative distances among response vectors (Fig. 2a; Kriegeskorte et al., 2008). Using RSA, a linear combination of features derived from different layers of a DNN has been shown to explain object representation in IT (Khaligh-Razavi and Kriegeskorte, 2014).

Figure 6a visualizes representational dissimilarity matrices (RDMs) for CMM and fMRI data for eight ROIs. Each element in the matrices indicates the normalized Euclidean distance, which measures the dissimilarity between activity patterns of different experimental conditions (Fig. 2a). We used the Kendall-𝛕_B_ correlation coefficient for comparing dissimilarity matrices, as indicated by the numbers between the two RDMs for each ROI.

**Figure 6.**
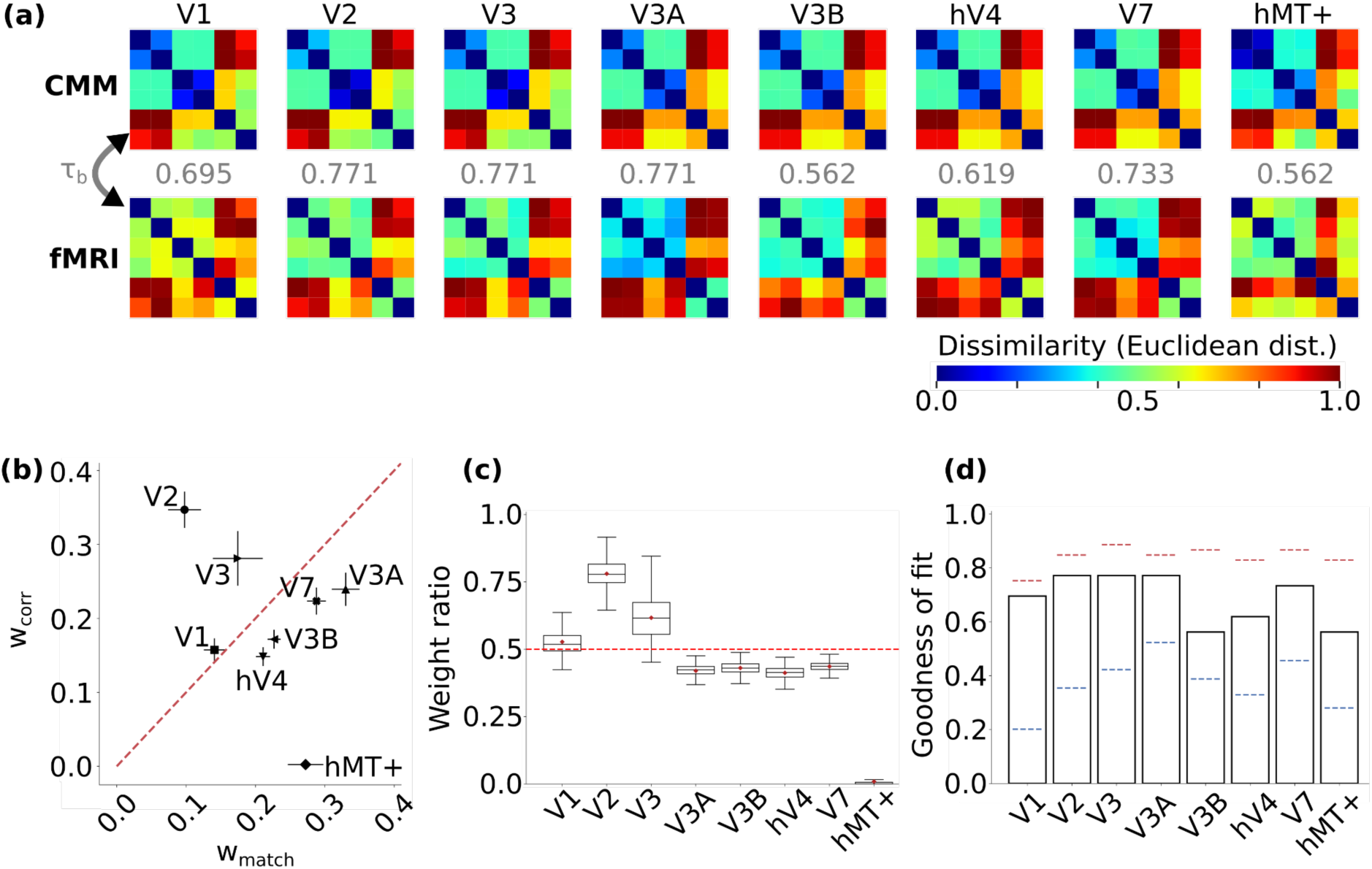
**(a)** Representational dissimilarity matrices (RDMs) for CMM and fMRI responses, using normalized Euclidean distance as the dissimilarity measure. The number below each RDM is the Kendall-𝛕_B_ correlation coefficient between the model and regional brain RDMs. **(b)** Proportions of cross-correlation (w_corr_) and cross-matching (w_match_) weights in each ROI, with error bars indicating bootstrap standard deviation (n = 1000). **(c)** Weight ratio index, w_corr_/(w_corr_ + w_match_), reflecting a preference for correlation representation. **(d)** Model’s goodness of fit, assessed by the Kendall-𝛕_B_ correlation coefficient. Blue and red dashed lines indicate the lower and upper bound of the noise ceiling, respectively. The model’s fit is within the noise ceiling, suggesting it accounts for the observed data variance.

The RDMs based on the CMM provided the best fit for the regional brain RDMs in V2, V3, and V3A (Kendall-𝛕_B_ = 0.771). In contrast, hMT+ had the lowest score (Kendall-𝛕_B_ = 0.562). By visual inspection, the similarity structure between CMM RDMs and regional brain RDMs in V1, V2, and V3 appear especially pronounced in between-category pairwise representational dissimilarities for aRDS-cRDS and aRDS-hmRDS clusters. However, CMM failed to capture within-category pairwise (crossed-uncrossed category within an RDS type) dissimilarity structure in aRDS-aRDS and hmRDS-hmRDS clusters. This suggests that the brain may adopt different computational strategies than those modeled by the CMM for representing within-category disparity in aRDSs and hmRDSs.

We estimated the contributions of cross-correlation (w_corr_) and cross-matching (w_match_) representations to regional brain representation using a non-negative least squares solution with an L2-norm applied to the linear fit between brain RDMs and CMM RDMs (Fig. 6b). To quantify the preference for the correlation-based representation, we calculated a weight ratio index w_corr_/(w_corr_ + w_match_), shown in Figure 6c.

Responses in early areas V1, V2, and V3 showed greater cross-correlation components (Figs. 6b, c); the weight ratio index was 0.528 in V1 (95% confidence interval (CI): 0.525 - 0.531), 0.780 in V2 (95% CI: 0.777 - 0.783), and 0.617 in V3 (95% CI: 0.612 - 0.622). These suggest that a majority of voxels in these areas responded primarily based on cross-correlation computation. Notably, V2 exhibited the highest weight ratio index, suggesting its prominent role in the correlation-based representation of binocular disparity.

In contrast, the weight ratio index values in V3A, V3B, V7, and hV4A were all below 0.5. Specifically, V3A showed a mean weight ratio index of 0.420 (95% CI: 0.418 - 0.422), V3B at 0.430 (95% CI: 0.428 - 0.432), V7 at 0.436 (95% CI: 0.435 - 0.437), and hV4 at 0.412 (95% CI: 0.410 - 0.414). These suggest a more dominant role of cross-matching mechanisms in processing disparity information in these areas. Most strikingly, the weight ratio value in hMT+ was close to 0, with a mean of 0.009 (95% CI: 0.007 - 0.011), signifying that disparity representation in this area can be almost perfectly explained by cross-matching computation.

The goodness of fit of CMM based on the Kendall-𝛕_B_ correlation coefficient fell between the lower and upper bounds of the noise ceiling (Fig. 6d). This indicates that the model RDMs could explain most of the variance observed in the brain RDMs.

Overall, the CMM analysis suggests that cross-correlation and cross-matching components were systematically distributed across the visual cortex. Disparity information in early areas up to V3 were strongly represented by cross-correlation, whereas cross-matching started to contribute with a greater weight after V3.

### GC-Net transforms stereo representation from cross-correlation to cross-matching across layers

The RDM analysis showed a transformation of disparity representation from cross-correlation dominance in early areas up to V3 to cross-matching dominance in areas anterior to V3. This raises two important questions. First, what is the driving force that makes this transformation happen? Second, how do individual neurons in each brain area process disparity from the two eyes to underlie this transformation?

We explored these questions by simulating disparity processing in a DNN optimized for processing binocular disparity. We employed GC-Net (Kendall et al., 2017) and trained it on the Scene Flow datasets consisting of three subsets (Monkaa, FlyingThings3D, and Driving subsets; Mayer et al., 2016) to estimate binocular disparity from stereo pairs of naturalistic images. Although we did not explicitly constrain the GC-Net to our RDSs or to our fMRI data, when trained on the Monkaa subset (8591 training frames with a resolution of 960 x 540 pixels), it exhibited depth judgment performance that was strikingly similar to that of humans: perfect for cRDSs, significantly above chance for hmRDSs, and below chance (i.e., reversed depth) for aRDSs (Fig. 1c, see Wundari and Ban, 2024). In contrast, training on the FlyingThings3D and Driving subsets did not yield human-like performance. Training on the FlyingThings3D results in a network that performed perfectly for both cRDSs and hmRDSs but failed to show reversed depth for aRDSs. Training on the Driving dataset produced a network with perfect performance for cRDSs but chance-level performance for both aRDSs and hmRDSs. Therefore, we used the GC-Net trained on the Monkaa subset for the following analysis.

We examined whether GC-Net undergoes a representational transformation from cross-correlation to cross-matching, as reflected in our empirical fMRI data and CMM simulations. We characterized the computational types of processing units in relation to cross-correlation and cross-matching in GC-Net layers by computing the AUC ratio (Fig. 2c; see Materials and Methods). An AUC ratio of zero indicates an ideal unit for cross-matching computation, and an AUC ratio of one indicates an ideal cross-correlation unit. The analysis began at layer 19, where the left and right input representations first interact.

Each GC-Net layer was populated by a different proportion of cross-correlation and cross-matching units. The AUC ratios for layers 19 and 20 predominantly fell within a narrow range between 0.6 and 0.8, indicating a strong influence of cross-correlation (Fig. 7a). The AUC ratios for layers 21-27 were distributed widely between zero and one, reflecting a mix of cross-correlation and cross-matching processes. Starting from layer 25, the AUC ratio began to bias towards values below 0.5. Specifically, nearly all units in layers 31-37 had AUC ratios below 0.5, indicating a greater contribution by cross-matching components in representing disparity. It should be noted that a small group of units with a greater cross-correlation component reappeared in layers 36 and 37. The AUC ratio analysis across GC-Net layers suggests that, similar to our fMRI analysis results, early layers constituted more units inclined toward cross-correlation than in later stages. Layers in the higher hierarchy were influenced more strongly by units of cross-matching nature but retained a small number of units of cross-correlation nature

**Figure 7.**
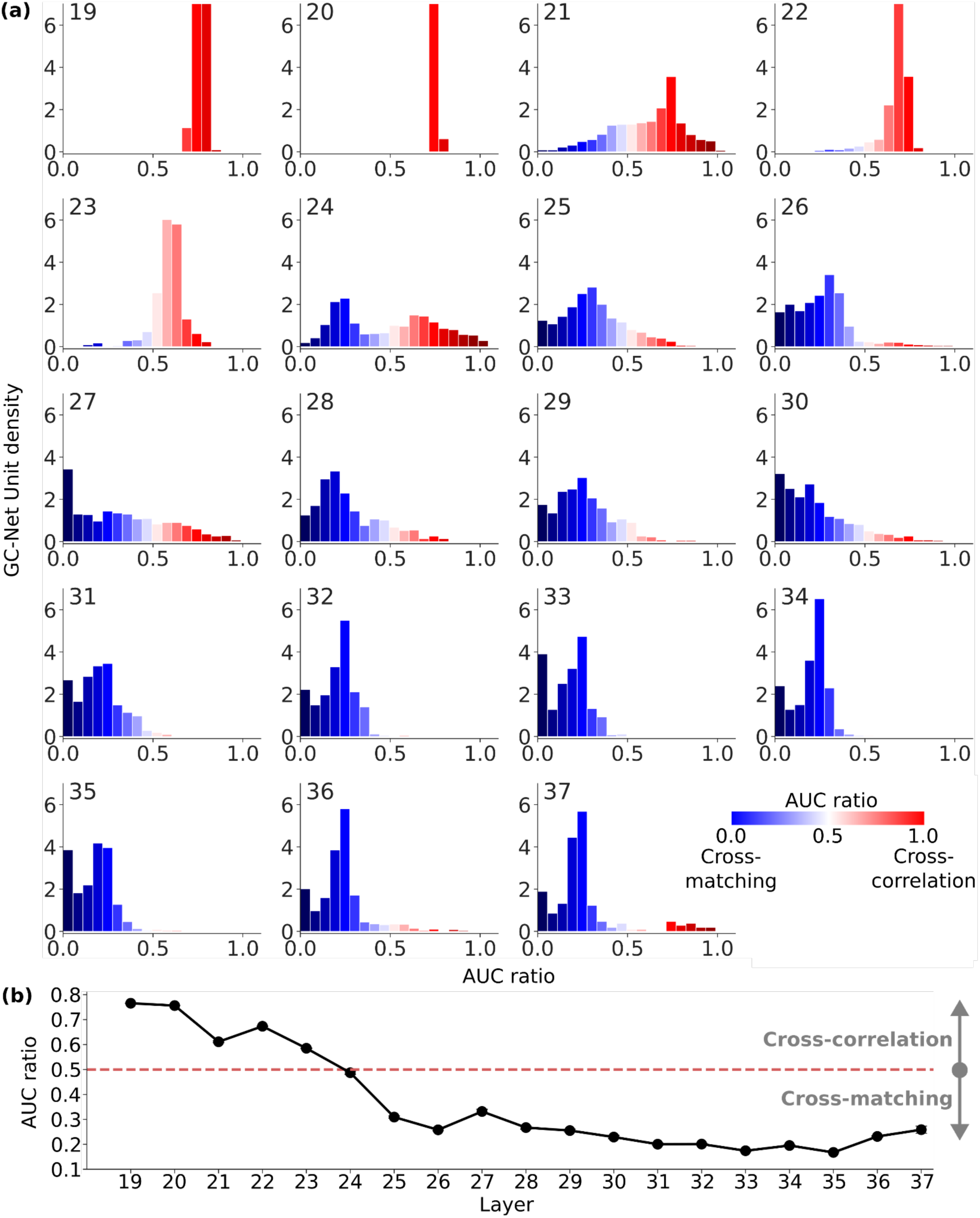
**(a)** Distribution of GC-Net units’ preferences for cross-correlation and cross-matching in each layer. The number on the top left indicates the layer name. **(b)** Evolution of cross-correlation and cross-matching preferences across DNN layers. This plot displays the mean AUC ratio calculated from the distributions in **(a)**, with error bars (barely visible) representing the standard error across units.

This trend is highlighted in plots of the mean AUC ratios across layers, depicting a clear transformation in disparity representations from cross-correlation to cross-matching across layers (Fig. 7b). Specifically, the mean of AUC ratios in layers 19-23 were above 0.5, suggesting a greater number of units engaged in cross-correlation. These layers, thus, share similarities with early visual areas V1-V3 in the human brain which showed a higher contribution from cross-correlation (Figs. 6b, c). The distribution of AUC ratio was bimodal in layer 24, and the mean AUC ratio was close to 0.5 (mean = 0.487, SE = 0.00674), indicating an equal portion of cross-correlation and cross-matching units. The majority of units in the subsequent layers (25-37) had AUC ratios below 0.5, indicating a greater number of units engaged in cross-matching. These layers show similarity with higher brain regions V3A/B, hV4, V7, and hMT+, which involve a greater cross-matching component (Figs. 6b, c).

Overall, the GC-Net analysis suggests two points. First, similar to the brains of monkeys and humans, an artificial visual system optimized for estimating binocular disparity exhibits representational transformation from cross-correlation to cross-matching. Second, the optimization for extracting stereoscopic depth in natural scenes may shape the neural mechanism for transforming disparity representation from cross-correlation to cross-matching.

### Characterization of the left and right preferred inputs across GC-Net layers

The theoretical disparity tuning of cross-correlation and cross-matching neurons (Fig. 1b) can arise from various configurations of binocular simple cells’ receptive fields (Read et al., 2002). However, the increasing complexity of disparity processing as one ascends the visual hierarchy poses challenges in estimating receptive fields from neural data. Consequently, we still lack a comprehensive understanding of how cross-correlation and cross-matching neurons in different visual areas integrate the two retinal inputs to support the transformation of disparity representation.

To address this issue, we characterized the left and right preferred inputs, or receptive fields, of GC-Net units across layers — what a GC-Net unit should “see” to maximize its activation. We estimated the respective preferred inputs using activation maximization (Zeiler and Fergus, 2014; Voss et al., 2021) and calculated Pearson’s correlation between them to assess their spatial structure similarity. We performed this analysis for the central units in each disparity and feature channel across layers 19-37.

We divided GC-Net layers 19-37 into three groups: early, middle, and late layers. The first six layers (19-24) were designated as early layers because they had a mean AUC ratio close to or greater than 0.5 (Fig. 7b), similar to early visual areas V1-V3, which had a mean weight ratio index greater than 0.5 (Fig. 6c). For the remaining 13 layers (25-37), we simply divided them into seven middle layers (25-31) and six late layers (32-37).

Figure 8a shows Pearson’s correlation coefficients between the left and right preferred inputs across layers 19-37. Each layer constituted a diverse range of units exhibiting interocular correlation from positive to negative, but with variations centered around different values (left panel). A negative correlation suggests that the units may respond to features that are present in one input channel or “eye”, but absent in the other. By plotting the mean AUC ratio against the mean Pearson’s correlation coefficient, the units in early, middle, and late layers were organized into three clusters (right panel).

**Figure 8.**
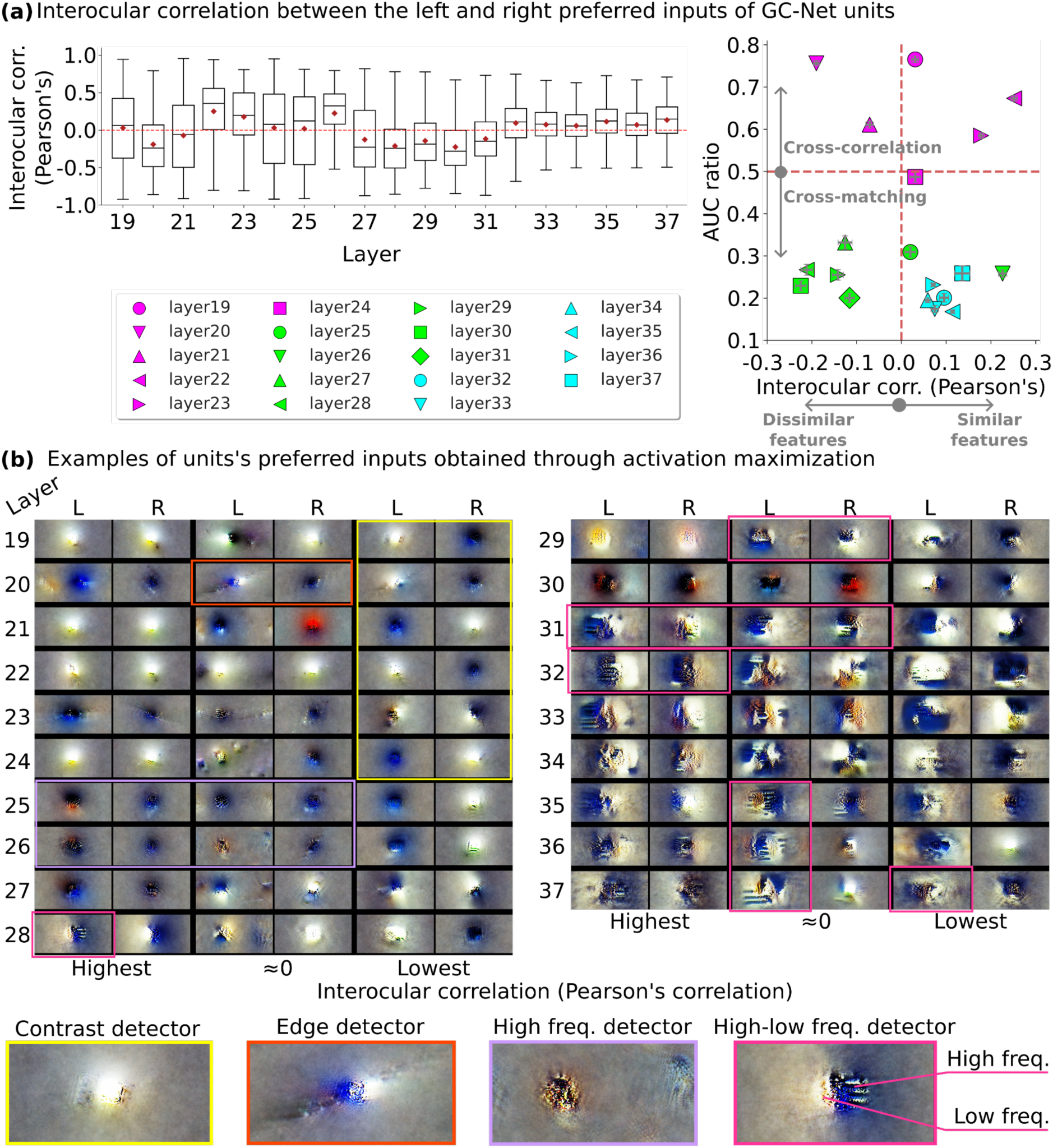
**(a)** Left: interocular correlation of preferred inputs for DNN units across layers, measured by Pearson’s correlation between the left and right preferred inputs obtained through activation maximization. Right: the mean Peason’s correlation coefficient plotted against the mean AUC ratio. Each cluster represents the early (magenta, 19-24), middle (green, 25-31), and late (cyan, 32-37) layers. The error bars (barely visible) represent standard errors across units. **(b)** Examples of units’ preferred inputs in RGB that maximize their activation (RFs). For each layer, three pairs are shown, corresponding to the highest, nearly zero, and lowest interocular correlation scores. Profiles delineated with colored rectangles are some detectors that can be recognized as contrast (yellow), edge (orange), high spatial frequency (purple), and high-low spatial frequency (pink) detectors.

The early stage of disparity processing (layers 19-24, magenta markers in Fig. 8a right panel) included a diverse set of units, some responding to single image features that are displaced by disparity, and others responding to different features in the two eyes. Specifically, layers 19, 22, 23, and 24 exhibited positive mean Pearson’s correlation coefficients [(mean, SE) = (0.031, 0.008), (0.252, 0.007), (0.178, 0.006), (0.031, 0.012), respectively], suggesting that, on average, units in these layers are strongly activated by similar image features that are displaced by disparity. In contrast, layers 20 and 21 showed negative mean coefficients [(mean, SE) = (−0.19, 0.006), (−0.071, 0.008), respectively], suggesting that these units, on average, respond maximally to false matches where image features in one eye do not correspond to features in the other. Such diversity suggests that the early stage of stereo processing involves a broad comparison of left and right inputs.

Nearly all preferred inputs for the early layers appeared Gaussian-shaped (examples shown in Fig. 8b). Preferred inputs with the highest correlation appeared identical. A unit profile with nearly zero correlation in layer 20 (delineated by an orange rectangle) shows one eye detecting a tilted edge structure and the other a circular structure of dark contrast. Preferred inputs for units with the lowest correlation in these early layers (delineated by a yellow rectangle) tended to have an opposite luminance, in slightly disparate positions between the eyes.

For the middle layers (green markers in Fig. 8a), five out of seven layers (27-31) had negatively correlated preferred inputs [mean, SE = (−0.125, 0.016), (−0.212, 0.013), (−0.142, 0.012), (−0.225, 0.012), (−0.116, 0.011), respectively]. This negative correlation suggests that a majority of cross-matching units as indicated by their AUC ratio below 0.5 in the middle layers achieve peak activity when detecting dissimilar features with an opposite contrast polarity.

Some profiles of the preferred inputs in the middle layers still resembled Gaussian shapes but with high spatial frequency structures inside (delineated by a light purple rectangle in Fig. 8b). The size of their spatial structure in layers 29-31 was larger than those in the early layers. We also observed an interesting profile in layer 28, delineated by a pink rectangle, in which a unit responded to low spatial frequency structures on one side of its RF, and high spatial frequency patterns on the other side (a high-low frequency detector). This detector may be useful for detecting the boundaries of objects when the background is out of focus (Olah et al., 2020b) or for detecting texture boundaries.

Late layers (32-37) exhibited AUC ratio below 0.5 and positive interocular correlation [(mean, SE) = (0.095, 0.01), (0.074, 0.008), (0.059, 0.006), (0.114, 0.004), (0.071, 0.004), (0.136, 0.017), respectively]. This suggests that cross-matching units in the late layers are strongly activated by features that appear similarly in both eyes. Features in these late layers appeared more complex and larger than those in the early layers. Some of them also show high-low frequency structures (delineated by pink rectangles in Fig. 8b). Other feature detectors in the deeper layers of GC-Net, however, cannot be straightforwardly interpreted. This might stem from the vastness of the image space, allowing for the generation of images that meet the objective function criteria yet remain unrecognizable (Nguyen et al., 2015).

## Discussion

We investigated how the human visual cortex represents binocular disparity by analyzing fMRI responses to disparities in visual stimuli with full match (cRDSs), false match (aRDSs), and half-match (hmRDSs) dot patterns between the eyes. In all ROIs, mean responses were higher for cRDSs compared to aRDSs and hmRDSs, with no significant difference between aRDSs and hmRDSs mean responses (Fig. 4). Nevertheless, disparity information in aRDSs was decodable from voxel response patterns in V3A and V7, while hmRDS disparity information was reliably detected in V2, V3, and V3A (Fig. 5). RSA revealed that V1, V2, and V3 mainly encoded disparity through cross-correlation, whereas V3A/B, V7, and hV4 relied more on cross-matching, and hMT+ strongly exhibited cross-matching (Fig. 6). A DNN optimized for disparity estimation showed a clear transformation from cross-correlation to cross-matching across layers (Fig. 7). Activation maximization suggests three phases of this transformation: initial processing of similar and dissimilar features from the two eyes, intermediate processing of dissimilar features, and final processing of similar features (Fig. 8).

### Distribution of cross-correlation and cross-matching representations across visual cortices

The three RDS types vary in dot-correlation and dot-matched levels: cRDSs (100% correlation, 100% matched), aRDSs (−100% correlation, 0% matched), and hmRDSs (0% correlation, 50% matched) (Doi et al., 2011). We leveraged this uniqueness to distinguish cross-correlation from cross-matching responses. Our participants experienced perceptual depth in cRDSs (veridical), hmRDSs (weaker veridical), and aRDSs (reversed) (Fig. 1c), consistent with Doi et al. (2011, 2013). Although mean BOLD responses in any visual areas examined could not explain the perceptual differences between hmRDSs and aRDSs (Figs. 1c, 4), MVPA revealed that their decoding accuracy varied (Fig. 5), suggesting diverse contributions of cross-correlation and cross-matching components across visual areas. RSA suggests a representational transformation from cross-correlation to slightly cross-matching up to middle stages (Fig. 6). It remains unclear how the transformation proceeds in higher stages, such as LOC in the temporal lobe and IPSs in the parietal lobe.

Preston et al. (2008) reported that disparity representation for anticorrelated features attenuated along the dorsal pathway V1-V2-V3-V3A-V7. However, our MVPA results showed increasingly robust disparity representation of aRDSs from V1 to V7, peaking at V3A (Fig. 5). The discrepancy might stem from the use of distinct RDSs between the two studies. Preston and colleagues presented static aRDSs for 500 ms, potentially triggering binocular rivalry, causing perceptual alternations and impairing stereopsis (Wolfe, 1983; Blake et al., 1991; Blake and Logothetis, 2002). This likely prevented their participants from perceiving depth in aRDSs, affecting the decoding accuracy due to the neural dynamics of interocular competition (Blake and Logothetis, 2002; Tong et al., 2006).

Conversely, the rapid flickering of dynamic aRDSs used in our study did not trigger binocular rivalry (Wolfe, 1983; O’Shea and Blake, 1986), allowing stereopsis to occur. Yet, V1 decoding accuracy for aRDSs remained at baseline (Fig. 5), likely resulting from the attenuation of neural activity corresponding to anticorrelated features (Cumming and Parker, 1997; Tanabe et al., 2011; Henrikson et al., 2016), which made the classifier fail at discriminating responses between near and far. Notably, in V3A, the classifier reliably decoded disparity information from aRDS response patterns, possibly due to its clustered organization of disparity preferences (Goncalves et al., 2015).

### Neural representations of false matches persist in the visual pathway

The transformation of disparity representation we demonstrated aligns with the traditional assumption that false match responses diminish along the visual hierarchy, as correct global matching requires rejecting false matches (Julesz, 1960; Marr and Poggio, 1976). However, this does not hold for every aspect of stereopsis. First, reversed depth perception for aRDSs provides strong evidence that the brain retains and utilizes false matches represented by cross-correlation for depth perception (Fujita and Doi, 2016). Second, in real 3D environments, such as occlusions, the brain exploits unmatched features for depth perception (Nakayama and Shimojo, 1990).

MVPA showed that false match information in aRDSs became increasingly robust from V1 to V3A (Fig. 5). Disparity representation carrying partial false matches (hmRDSs) was also enhanced from V1 to V3A. In contrast, disparity information in aRDSs and hmRDSs attenuated in hV4, contrary to Preston et al. (2008), who reported robust aRDS decoding accuracy in hV4, comparable to cRDS. These suggest that false match information is incorporated, rather than attenuated or discarded, in the early stages of the dorsal pathway. V3A has a larger population RF than V1 (Alvares et al., 2021), allowing for more effective contextual processing. Thus, V3A may detect global matching levels, aiding other brain regions in depth perception, object recognition, and navigation.

### GC-Net exhibits a transformation from cross-correlation to cross-matching

CMM, assuming two parallel representations for depth (Figs. 1b and 2a), demonstrated a progression from cross-correlation to slightly cross-matching up to mid-stages (Fig. 6). GC- Net, which assumed a single hierarchical processing stream and did not use fMRI data in the optimization, also showed unit responses’ transformation from cross-correlation to cross-matching across its layers (Fig. 7). Depth maps generated by GC-Net were encoded by cross-correlation and cross-matching units in the final layer, operating similarly to CMM. Remarkably, GC-Net’s depth judgment matched human performance (Fig. 1c). These suggest that optimization for depth extraction shapes the neural mechanism for stereo processing in the brain, similar to the principle underlying visual object recognition (Yamins and DiCarlo, 2016; Yamins et al., 2014).

Analysis of the units’ preferred inputs suggests three phases occurring during the transformation for solving the correspondence problem: an early phase extracting local binocular information from similar and dissimilar features, a middle phase focusing on dissimilarity processing, and a final phase emphasizing matching processing (Fig. 8a).

In the early layers, small Gaussian-like filters process disparity by extracting local binocular information from similar and dissimilar features (Fig. 8). V1, V2, and V3 fit this role, encoding disparity with a focus on cross-correlation sensitive to both similar and dissimilar features (Figs. 6b, c). This mechanism resonates with disparity-selective neurons in monkey V1, many of which exhibit hybrids of position-shift and phase-shift of monocular RFs (Anzai et al., 1999; Prince et al., 2002), useful for detecting false matches (Read and Cumming 2007; Goncalves and Welchman 2017). V2 neurons sensitive to depth stratification or figure-ground segregation (von der Heydt et al., 2000; Qiu and von der Heydt, 2005, 2007) may require processing dissimilar features between the eyes.

The middle stages shift towards processing dissimilarity, indicated by their mean negative interocular RF correlation. This suggests that the transformation may benefit from processing dissimilar features, possibly by driving the suppressive units’ activity to provide more information about unlikely interpretations of the scene (proscription strategy, Tanabe et al., 2011; Goncalves and Welchman, 2017). Comparing the AUC ratio (Fig. 8b) and the cross-correlation preference (Fig. 7c), we propose that this intermediate attribute strongly resides in mid-brain areas V3A/B, V7, and hV4, supporting reversed depth perception without accompanying robust shape perception (Tanabe et al., 2008).

In the final stage, disparity processing focuses on similarity, suggesting that cross-matching features emerge after the early and middle stages partially solve the correspondence problems. This finding is consistent with the findings in monkey IT and AIP, where neurons do not respond to anticorrelated features (Janssen et al., 2003; Theys et al., 2012).

### Comparison with neurophysiology studies in non-human primates

Monkey electrophysiology showed that V1 neurons extract binocular disparity using cross-correlation (Cumming and Parker, 1997), while higher cortices, V4, IT, and AIP, rely on cross-matching (Janssen et al., 2003; Tanabe et al., 2004; Theys et al., 2012; Abdolrahmani et al., 2016; Yoshioka et al., 2021). Similarly, our results showed that disparity representation in early human visual areas V1, V2, and V3 was closer to cross-correlation. Higher visual areas V3A/B, V7, and hV4 shifted slightly towards cross-matching.

Our finding that hMT+ strongly favored cross-matching (Fig. 6) aligns with prior human studies (Bridge and Parker 2007; Preston et al., 2008), but contrasts sharply with monkey studies (Krug et al., 2004; Yoshioka et al., 2021), where MT responses are better explained by cross-correlation. The discrepancy might arise from several possibilities. First, hMT+ encompasses multiple areas besides MT, including a region homologous to the monkey MST (Huk et al., 2002). Moreover, retinotopic mapping delineates visual areas in this middle temporal region differently between human and monkey cortices and even within humans depending on the definition (Amano et al., 2009; Huk et al., 2012; Kolster et al., 2010, 2014). Second, while hMT+ neurons are in fact sensitive to disparity in aRDSs, variability in the phase shift of tuning across nearby neurons could average out disparity-sensitive responses to aRDSs due to the spatial pooling of BOLD signals (Doi et al., 2018). Lastly, the difference may reflect an evolutionary divergence between species, possibly driven by distinct ecological and environmental pressures (Armendariz et al., 2019).

To conclude, we show that disparity representation is systematically distributed between the early and higher visual cortex in which cross-matching plays a greater role in regions anterior to V3. Driven by optimization for extracting depth in natural scenes, disparity representation transforms from cross-correlation to cross-matching along the visual pathways. Disparity encoding from dissimilar features may be crucial in facilitating the transformation.

## Acknowledgments

This research was supported by a JST ERATO grant to H.B. (JPMJER1801) and grants to I.F. (17H04790 and 21H02596) and H.B. (21H00968 and 21K18572) from the Ministry of Education, Culture, Science, Sports and Technology. We thank Drs. Kaoru Amano, Dorita Chang, Takahiro Doi, and Hiromasa Takemura for their comments on earlier versions of the manuscript.

## Author contributions

B.G.W, I.F., and H.B. designed research; B.G.W., and H.B. performed research; B.G.W., and H.B. analyzed data; B.G.W., I.F., and H.B. wrote the paper.

## References

Abdolrahmani M, Doi T, Shiozaki HM, Fujita I (2016) Pooled, but not single-neuron, responses in macaque v4 represent a solution to the stereo correspondence problem. J Neurophysiol 115:1917–1931.

Adams DL, Zeki S (2001) Functional organization of macaque v3 for stereoscopic depth. J Neurophysiol 86:2195–2203.

Alvarez I, Hurley SA, Parker AJ, Bridge H (2021) Human primary visual cortex shows larger population receptive fields for binocular disparity-defined stimuli. Brain Structure and Function 226:2819–2838.

Amano K, Wandell BA, Dumoulin SO (2009) Visual field maps, population receptive field sizes, and visual field coverage in the human MT+ complex. J Neurophysiol 102:2704–2718.

Anzai A, Ohzawa I, Freeman RD (1999) Neural mechanisms for encoding binocular disparity: receptive field position versus phase. J Neurophysiol 82:874–890.

Aoki SC, Shiozaki HM, Fujita I (2017) A relative frame of reference underlies reversed depth perception in anticorrelated random-dot stereograms. J Vis 17:17–17.

Armendariz M, Ban H, Welchman AE, Vanduffel W (2019) Areal differences in depth cue integration between monkey and human. PLOS Biol 17:e2006405.

Ban H, Preston TJ, Meeson A, Welchman AE (2012) The integration of motion and disparity cues to depth in dorsal visual cortex. Nat Neurosci 15:636–643.

Ban, H., Yamamoto, H. (2013) A non-device-specific approach to display characterization based on linear, nonlinear, and hybrid search algorithms. J Vision, 13(6):20, 1-26.

Blake R, Logothetis NK (2002) Visual competition. Nature Rev Neurosci 3:13–21.

Blake R, Yang YD, Wilson HR (1991) On the coexistence of stereopsis and binocular rivalry. Vision Res 31:1191–1203.

Bridge H, Parker AJ (2007) Topographical representation of binocular depth in the human visual cortex using fMRI. J Vis 7:15.1–1514.

Chen G, Lu HD, Roe AW (2008) A map for horizontal disparity in macaque V2. Neuron 58:442–450.

Cumming BG, Parker AJ (1997) Responses of primary visual cortical neurons to binocular disparity without depth perception. Nature 389:280–283.

DeAngelis GC, Newsome WT (1999) Organization of disparity-selective neurons in macaque area MT J Neurosci 19:1398–1415.

DeYoe EA, Carman GJ, Bandettini P, Glickman S, Wieser J, Cox R, Miller D, Neitz J (1996) Mapping striate and extrastriate visual areas in human cerebral cortex. Proc Natl Acad Sci USA 93:2382–2386.

Doi T, Abdolrahmani M, Fujita I (2018) Spatial pooling inherent to intrinsic signal optical imaging might cause V2 to resemble a solution to the stereo correspondence problem. Proc Natl Acad Sci USA 115:E6967–E6968.

Doi T, Fujita I (2014) Cross-matching: a modified cross-correlation underlying threshold energy model and match-based depth perception. Front Comput Neurosci 8:127, 1–15.

Doi T, Takano M, Fujita I (2013) Temporal channels and disparity representations in stereoscopic depth perception. J Vis 13:26–26.

Doi T, Tanabe S, Fujita I (2011) Matching and correlation computations in stereoscopic depth perception. J Vis 11:1–1.

Dumoulin SO, Wandell BA (2008) Population receptive field estimates in human visual cortex. Neuroimage 39:647–660.

Erhan D, Bengio Y, Courville A, Vincent P (2009) Visualizing higher-layer features of a deep network. Tech Rep Univ Montréal 1341:1–13.

Etzel JA (2015) MVPA permutation schemes: permutation testing for the group level. In Proceeding of 2015 International Workshop on Pattern Recognition in NeuroImaging, IEEE pp. 65–68.

Fujita I, Doi T (2016) Weighted parallel contributions of binocular correlation and match signals to conscious perception of depth. Philos Trans R Soc Lond B Biol Sci 371:20150257.

Goncalves NR, Ban H, Sánchez-Panchuelo RM, Francis ST, Schluppeck D, Welchman AE (2015) 7 tesla fMRI reveals systematic functional organization for binocular disparity in dorsal visual cortex. J Neurosci 35:3056–3072.

Goncalves NR, Welchman AE (2017) “What not” detectors help the brain see in depth. Curr Biol 27:1403–1412.

Güçlü U, van Gerven MA (2015) Deep neural networks reveal a gradient in the complexity of neural representations across the ventral stream. J Neurosci 35:10005–10014.

Henriksen S, Read JC, Cumming BG (2016) Neurons in striate cortex signal disparity in half-matched random-dot stereograms. J Neurosci 36:8967–8976.

Huk AC, Dougherty RF, Heeger DJ (2002) Retinotopy and functional subdivision of human areas MT and MST. J Neurosci 22:7195–7205.

Janssen P, Vogels R, Liu Y, Orban GA (2003) At least at the level of inferior temporal cortex, the stereo correspondence problem is solved. Neuron 37:693–701.

Julesz B (1960) Binocular depth perception of computer-generated patterns. Bell Syst Tech J 39:1125–1162.

Kamitani Y, Tong F (2005) Decoding the visual and subjective contents of the human brain. Nat Neurosci 8:679–685.

Kendall A, Martirosyan H, Dasgupta S, Henry P, Kennedy R, Bachrach A, Bry A (2017) End-to-end learning of geometry and context for deep stereo regression In Proc IEEE Int Conf Comput Vis, pp. 66–75.

Khaligh-Razavi SM, Kriegeskorte N (2014) Deep supervised, but not unsupervised, models may explain IT cortical representation. PLOS Comput Biol 10:e1003915.

Kleiner M, Brainard D, Pelli D, Ingling A, Murray R, Boussard C (2007) What’s new in psych toolbox-3? Perception 36:1.

Kolster H, Janssens T, Orban GA (2014) The retinotopic organization of macaque occipitotemporal cortex anterior to V4 and caudoventral to the middle temporal (MT) cluster. J Neurosci 34:10168–1069.

Kolster H, Peeters R, Orban GA (2010) The retinotopic organization of the human middle temporal area MT/V5 and its cortical neighbors. J Neurosci 30:9801–9820.

Kriegeskorte N (2011) Pattern-information analysis: from stimulus decoding to computational-model testing. Neuroimage 56:411–421.

Kriegeskorte N (2015) Deep neural networks: a new framework for modeling biological vision and brain information processing. Annu Rev Vis Sci 1:417–446.

Kriegeskorte N, Diedrichsen J (2016) Inferring brain-computational mechanisms with models of activity measurements. Philos Trans R Soc Lond B Biol Sci 371:20160278.

Kriegeskorte N, Goebel R, Bandettini P (2006) Information-based functional brain mapping. Proc Natl Acad Sci USA 103:3863–3868.

Kriegeskorte N, Mur M, Bandettini PA (2008) Representational similarity analysis-connecting the branches of systems neuroscience. Front in syst neurosci 2:249.

Kriegeskorte N, Wei XX (2021) Neural tuning and representational geometry. Nat Rev Neurosci 22:703–718.

Krug K, Cumming BG, Parker AJ (2004) Comparing perceptual signals of single V5/MT neurons in two binocular depth tasks. J Neurophysiol 92:1586–1596.

Loshchilov I, Hutter F (2017) Decoupled weight decay regularization. arXiv:1711.05101 .

Marr D, Poggio T (1976) Cooperative computation of stereo disparity. Science 194:283–287.

Mayer N, Ilg E, Häusser P, Fischer P, Cremers D, Dosovitskiy A, Brox T (2016) A large dataset to train convolutional networks for disparity, optical flow, and scene flow estimation In IEEE Int Conf on Comp Vis and Pat Rec (CVPR) arXiv:1512.02134.

Nakayama K, Shimojo S (1990) Da Vinci stereopsis: depth and subjective occluding contours from unpaired image points. Vision Res 30:1811–1825.

Nguyen A, Yosinski J, Clune J (2015) Deep neural networks are easily fooled: High confidence predictions for unrecognizable images. In Proc of the IEEE conf on comp vis and pat rec, pp. 427–436.

Nichols TE, Holmes AP (2002) Nonparametric permutation tests for functional neuroimaging: a primer with examples. Human Brain Mapping 15:1–25.

Nieder A, Wagner H (2000) Horizontal disparity tuning of neurons in the visual forebrain of the behaving barn owl. J Neurophysiol 83:2967–2979.

Nili H, Wingfield C, Walther A, Su L, Marslen-Wilson W, Kriegeskorte N (2014) A toolbox for representational similarity analysis. PLOS Comput Biol 10:e1003553.

Ohzawa I, DeAngelis GC, Freeman RD (1990) Stereoscopic depth discrimination in the visual cortex: neurons ideally suited as disparity detectors. Science 249:1037–1041.

Olah C, Cammarata N, Schubert L, Goh G, Petrov M, Carter S (2020a) An overview of early vision in inceptionv1. Distill https://distill.pub/2020/circuits/early-vision.

Olah C, Cammarata N, Schubert L, Goh G, Petrov M, Carter S (2020b) Zoom in: An introduction to circuits. Distill https://distill.pub/2020/circuits/zoom-in.

O’Shea RP, Blake R (1986) Dichoptic temporal frequency differences do not lead to binocular rivalry. Percept Psychophys 39:59–63.

Paszke A et al. (2019) Pytorch: an imperative style, high-performance deep learning library. Adv Neural Inf Process Syst 32.

Popple AV, Smallman HS, Findlay JM (1998) The area of spatial integration for initial horizontal disparity vergence. Vision Res 38:319–326.

Preston TJ, Li S, Kourtzi Z, Welchman AE (2008) Multivoxel pattern selectivity for perceptually relevant binocular disparities in the human brain. J Neurosci 28:11315–11327.

Prince SJ, Cumming BG, Parker AJ (2002) Range and mechanism of encoding of horizontal disparity in macaque V1. J Neurophysiol 87:209–221.

Qiu FT, von der Heydt R (2005) Figure and ground in the visual cortex: V2 combines stereoscopic cues with gestalt rules. Neuron 47:155–166.

Qiu FT, von der Heydt R (2007) Neural representation of transparent overlay. Nat Neurosci 10:283–284.

Read JC (2023) Stereopsis without correspondence. Philos Trans R Soc B 378:20210449.

Read JC, Cumming BG (2007) Sensors for impossible stimuli may solve the stereo correspondence problem. Nat Neurosci 10:1322–1328.

Read JC, Parker AJ, Cumming BG (2002) A simple model accounts for the response of disparity-tuned V1 neurons to anticorrelated images. Vis Neurosci 19:735–753.

Rosenberg A, Thompson LW, Doudlar R, Chang T-Y (2023) Neuronal representations supporting three-dimensional vision in nonhuman primates. Annu Rev Vis Sci 9:5.1– 5.23.

Sereno MI, Dale AM, Reppas JB, Kwong KK, Belliveau JW, Brady TJ, Rosen BR, Tootell RB (1995) Borders of multiple visual areas in humans revealed by functional magnetic resonance imaging. Science 268:889–893.recognize

Swisher JD, Halko MA, Merabet LB, MaMains SA, Somers DC (2007) Visual topography of human intraparietal sulcus. J Neurosci 27:5326–5337.

Tanabe S, Doi T, Umeda K, Fujita I (2005) Disparity-tuning characteristics of neuronal responses to dynamic random-dot stereograms in macaque visual area V4. J Neurophysiol 94:2683–2699.

Tanabe S, Haefner RM, Cumming BG (2011) Suppressive mechanisms in monkey v1 help to solve the stereo correspondence problem. J Neurosci 31:8295–8305.

Tanabe S, Umeda K, Fujita I (2004) Rejection of false matches for binocular correspondence in macaque visual cortical area v4. J Neurosci 24:8170–8180.

Tanabe S, Yasuoka S, Fujita I (2008) Disparity-energy signals in perceived stereoscopic depth. J Vis 8:22.1–2210.

Theys T, Srivastava S, van Loon J, Goffin J, Janssen P (2012) Selectivity for three-dimensional contours and surfaces in the anterior intraparietal area. J Neurophysiol 107:995–1008.

Tong F, Meng M, Blake R (2006) Neural bases of binocular rivalry. Trends in cognitive sciences 10:502–511.

von der Heydt R, Zhou H, Friedman HS (2000) Representation of stereoscopic edges in monkey visual cortex. Vision Res 40:1955–1967.

Voss C, Cammarata N, Goh G, Petrov M, Schubert L, Egan B, Lim SK, Olah C (2021) Visualizing weights. Distill https://distill.pub/2020/circuits/visualizing-weights.

Welchman AE (2016) The human brain in depth: how we see in 3D. Annu Rev Vis Sci 2:345– 376.

Wolfe JM (1983) Influence of spatial frequency, luminance, and duration on binocular rivalry and abnormal fusion of briefly presented dichoptic stimuli. Perception 12:447–456.

Wundari BG, Ban H (2024) Reversed depth representation in human and artificial visual systems. In Vis Sci Soc–VSS 2024: 24th Annual meeting of the Vision Sciences Society, Florida, US, May 17-22, 2024. VSS.

Yamamoto H, Ban H, Fukunaga M, Tanaka C, Umeda M, Ejima Y (2008) Large- and small-scale functional organization of visual field representation in the human visual cortex. In Visual cortex: new Research (eds. T. A. Portocello and R. B. Velloti) pp. 195–226. New York: Nova Science Publishers.

Yamins DL, DiCarlo JJ (2016) Using goal-driven deep learning models to understand sensory cortex. Nat Neurosci 19:356–365.

Yamins DL, Hong H, Cadieu CF, Solomon EA, Seibert D, DiCarlo JJ (2014) Performance-optimized hierarchical models predict neural responses in higher visual cortex. Proc Natl Acad Sci U S A 111:8619–8624.

Yoshioka TW, Doi T, Abdolrahmani M, Fujita I (2021) Specialized contributions of mid-tier stages of dorsal and ventral pathways to stereoscopic processing in macaque. eLife 10:e58749.

Yoshiyama K, Uka T, Tanaka H, Fujita I (2004) Architecture of binocular disparity processing in monkey inferior temporal cortex. Neurosci Res 48:155–167

Zeiler MD, Fergus R (2014) Visualizing and understanding convolutional networks in computer vision. ECCV 2014: 13th European Conference, Zurich, Switzerland, September 6–12, 2014, Proceedings, Part I 13, pp. 818–833. Springer.

Zhaoping L, Ackerman J (2018) Reversed depth in anti-correlated random dot stereograms and the central-peripheral difference in visual inference. Perception 47:531–539.

